# Measurement of selective constraint on human gene expression

**DOI:** 10.1101/345801

**Authors:** Emily C Glassberg, Ziyue Gao, Arbel Harpak, Xun Lant, Jonathan K Pritchard

## Abstract

Gene expression variation is a major contributor to phenotypic variation in human complex traits. Selection on complex traits may therefore be reflected in constraint on gene expression levels. Here, we explore the effects of stabilizing selection on *cis*-regulatory genetic variation in humans. We analyze patterns of expression variation at copy number variants and find evidence for selection against large increases in gene expression. Using allele-specific expression (ASE) data, we further show evidence of selection against smaller-effect variants. We estimate that, across all genes, singletons in a sample of 122 individuals have approximately 2.5 × greater effects on expression variance than common variants. Despite their increased effect sizes relative to common variants, we estimate that singletons in the sample studied explain, on average, only 5% of the heritability of gene expression from *cis*-regulatory variants. Finally, we show that genes depleted for loss-of-function variants are also depleted for *cis*-eQTLs and have low levels of allelic imbalance, confirming tighter constraint on the expression levels of these genes. We conclude that constraint on gene expression is present, but has relatively weak effects on most *cis*-regulatory variants, thus permitting high levels of gene-regulatory genetic variation.

## Introduction

Variation in human complex traits is connected to variation in gene expression levels (1–7). Selection on complex traits (i.e. complex disease) may therefore be reflected in selection on gene expression.

Across species, gene expression levels have been shown to evolve more slowly than would be expected under a neutral model (8–11). An analysis of mammalian gene duplications also showed that the total expression of gene pairs in species that experienced a duplication event is similar to the expression of the corresponding single gene copy in species without duplication (12). As a duplication event would be expected to dramatically increase gene expression, this observation suggests that stabilizing selection on gene expression may act to return the total expression of duplicate gene pairs to an optimal expression level.

Constraint on gene expression levels has also been shown to influence patterns of genetic variation within humans. First, some genes are unusually depleted for loss-of-function and copy number variants (13, 14). These genes are thought to be particularly constrained with respect to their expression.

Further, individuals with extreme expression levels for a particular gene are more likely to have rare variants in *cis* than individuals with average expression levels (15–18), suggesting that large gene expression changes are associated with rare genetic variation. As detailed below, this observation is consistent with stabilizing selection on expression levels selecting against large-effect regulatory variants.

Despite this evidence for constraint on expression levels, humans exhibit substantial variation in gene expression and possess many common gene regulatory variants (19–21). It is therefore of interest to quantify the breadth of constraint on gene expression levels across the human genome and its effects on gene-regulatory variation.

Under a model of stabilizing selection, the negative fitness effect of a *cis*-regulatory variant increases with its effect on expression level. In other words, as large-effect variants move individuals further from an optimal gene expression value, selection will act against them, keeping them more rare than those with small effects or no effect on expression. This would reduce the variance in gene expression levels and create a global relationship between the allele frequency and effect size of gene-regulatory variants (Fig 1; see Simons et al. 22 for a detailed model of stabilizing selection, genetic variation, and complex trait variance).

**Fig. 1.**
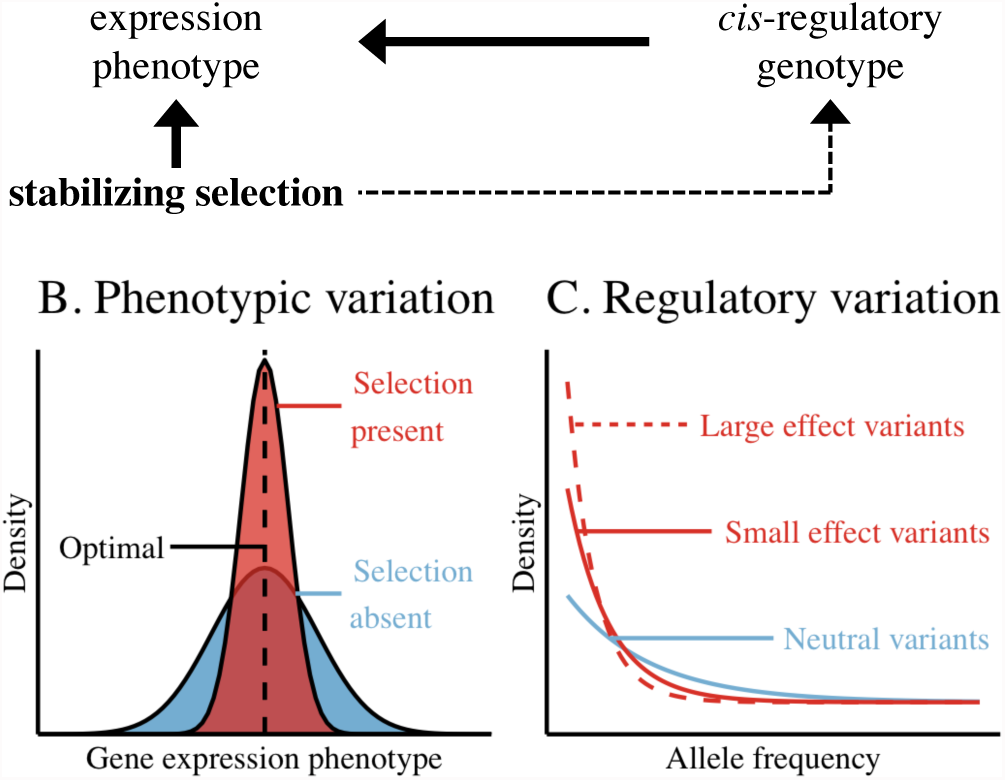
Expected signals of stabilizing selection on gene expression.

This model of stabilizing selection is consistent with observed patterns of variation of gene-regulatory variants. Past studies have noted a relationship between the allele frequency and effect size of expression quantitative trait loci (eQTLs) (20, 23). More recently, polygenic models have been shown to demonstrate a relationship between allele frequency and the variance in gene expression explained by trait-associated variants (24, 25). However, the strength and breadth of this selection across genes have yet to be quantified.

Here, we test whether patterns of genetic variation at gene-regulatory loci are consistent with human gene expression evolving under stabilizing selection. We analyze gene expression and regulatory genetic variation related to copy number variants (CNVs), expression quantitative trait loci (eQTLs), and allele-specifically expressed (ASE) transcripts. Together, these data show that, although constraint on expression affects patterns of gene expression variation and genetic variation at gene regulatory loci, its effects are relatively weak.

## Results

### Copy number variants show dosage sensitivity

Gene duplications and deletions are likely to have large effects on gene expression. If gene expression levels are under stabilizing selection, we would expect selection to keep large-effect copy number variants (CNVs) rare in the population. Therefore, to test whether human genetic variation is affected by selection against large changes in gene expression, we analyzed the expression levels and allele frequencies of CNVs. Using whole-genome and RNA sequencing of 147 individuals from version 4 of the GTEx Consortium (21), we identified 694 genes whose entire coding sequence had been duplicated in 196 polymorphic CNVs and obtained their expression levels across 12 tissues.

Despite high variance in expression ratios across genes, when a duplication CNV is rare (present in one individual in this sample), genes in the CNV are expressed at higher levels in heterozygous carriers (with three gene copies) than in non-carriers (with two gene copies; median expression ratio 1.31; Fig 2). By contrast, genes in common CNVs (>5% of sampled individuals have a duplicate gene copy) are expressed at similar levels in individuals with two and three copies (median expression ratio is 0.95; Fig 2).

**Fig. 2.**
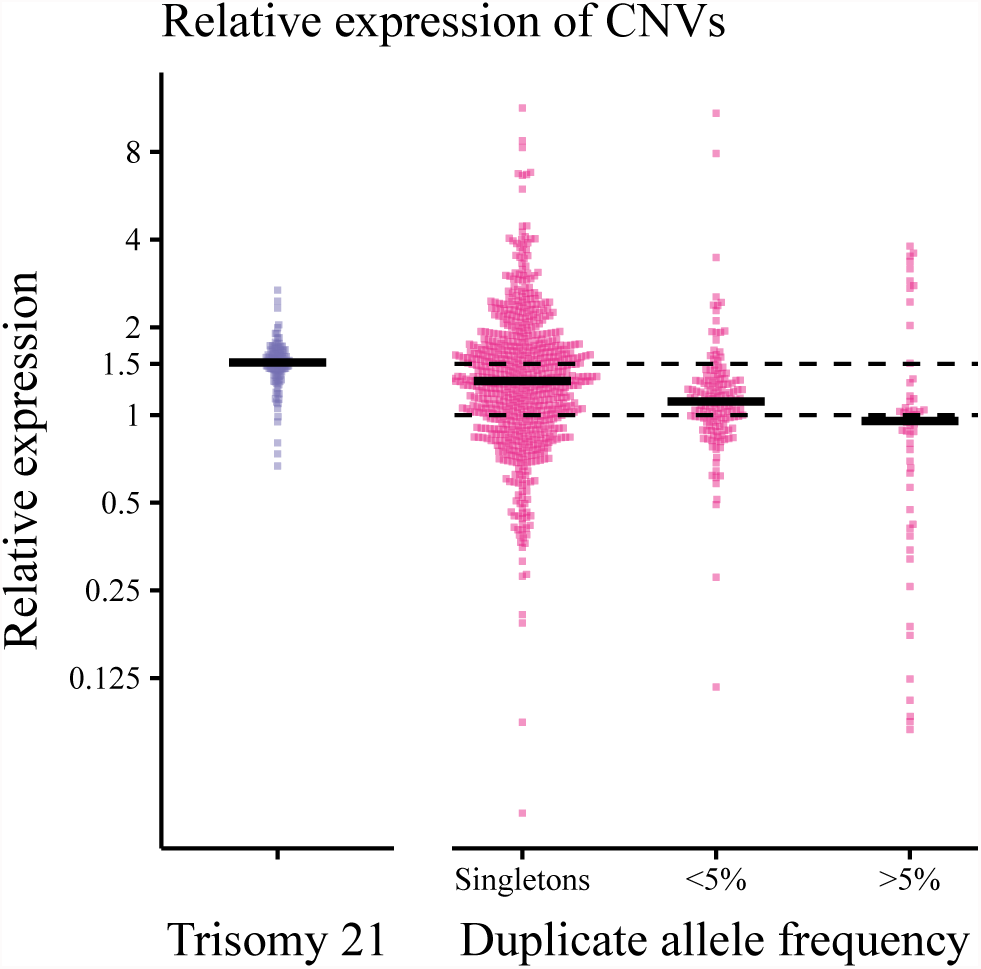
Expression of heterozygous carriers of a gene duplication (with three gene copies) relative to non-carriers (with two gene copies). On the left, each point corresponds to the expression level (RPKM) of one of 104 genes in an individual with trisomy 21 relative to the expression of the gene in their diploid twin. On the right, each point corresponds to the median expression of heterozygous duplication carriers relative to the median expression of non-carriers for one gene in one tissue. To reduce noise in the expression ratio, gene/tissue pairs with median non-carrier RPKM<1 were excluded (891 gene/tissue pairs retained). Duplicate frequencies are from GTEx. Black lines show median expression ratios across genes in each bin.

If each gene copy were expressed at the same, baseline level, one would expect the ratio of heterozygous duplication carriers to non-carriers to be 1.5. It is possible that observed expression ratio lower than 1.5 results from a buffering mechanism at the cellular level that returns the total expression level of each gene closer to that in the absence of the duplication. To test for expression buffering, we obtained gene expression measurements based on RNA sequencing data from a set of monozygotic twins discordant for trisomy 21 (26). Importantly, as trisomy 21 is a de novo expression-altering event, selection cannot affect the newly introduced expression changes.

We see that the average expression of genes on chromosome 21 is 50% higher in the trisomy 21 individual than in their diploid twin (Fig 2). This suggests that, before selection has time to act, when the entire gene and *cis*-regulatory region are duplicated, cellular buffering is negligible and expression levels increase proportionally with gene dosage. This is consistent with previous work showing that, for common, multi-allelic CNVs, gene expression levels scale linearly with copy number (27).

The lack of expression buffering after de novo chromosome duplication further supports the negative relationship between total expression level of duplicated genes and CNV frequency/age, which suggests constraint on gene expression levels. In other words, gene duplications can become common only when their total expression levels are comparable to the original levels.

Three possible mechanisms could explain this pattern. First, CNVs may arise on genetic backgrounds that vary in their expression levels. In this case, duplicates that arise on low-expression backgrounds (e.g., haplotypes with an expression-decreasing eQTL) are more likely to spread in the population. Second, during gene duplication, damage may occur to *cis*-regulatory elements of the duplicated gene. Damaged duplicates may lead to smaller expression increases and are therefore more likely to survive. Third, haplotypes with a duplicate gene may acquire additional genetic variation; haplotypes on which compensatory, down-regulating variants arise can become common.

Regardless of the mechanism, these patterns of expression variation at human CNVs confirm that constraint on gene expression has pronounced effects on genetic variants that cause large changes in expression.

However, many genetic variants may have smaller effects on expression than a gene duplication. We therefore investigated whether selection on gene expression is sufficiently strong to respond to subtler perturbations in expression levels.

### Characterizing patterns of *cis*-regulatory variation using expression quantitative trait loci

To detect genetic variants that affect gene expression, we first carried out expression quantitative trait locus (eQTL)-mapping to genotype and whole-blood RNA sequencing data from 922 European individuals from the Depression Genes and Networks cohort (20). To allow multiple independent causal variants per region, we used a stepwise regression approach to call eQTLs in 100kb windows centered on the transcription start site (TSS) of 12,794 autosomal, protein-coding genes (see). In total, we tested 3,309,888 SNPs and called 6,587 significant eQTLs associated with 4,734 genes (37% of genes tested).

To preserve interpretability of our effect size estimates, we did not perform any principal component or latent factor correction on our genotype or phenotype data, nor did we include any additional covariates in our eQTL mapping. As these additions would have improved our statistical power, the approximately two-fold fewer eGenes (genes with a significant *cis*-eQTL) detected here relative to previous eQTL studies of this dataset is expected (20).

Of genes with at least one significant *cis*-eQTL, 27.33% had more than one significant association (to a maximum of 9 eQTLs per gene). This suggests that there may, in many cases, be multiple, independent regulatory variants that would be missed in analyses of ‘lead eQTLs’ that consider only the most significant association per locus. Two example loci with multiple *cis*-eQTLs are shown in Fig 3.

**Fig. 3.**
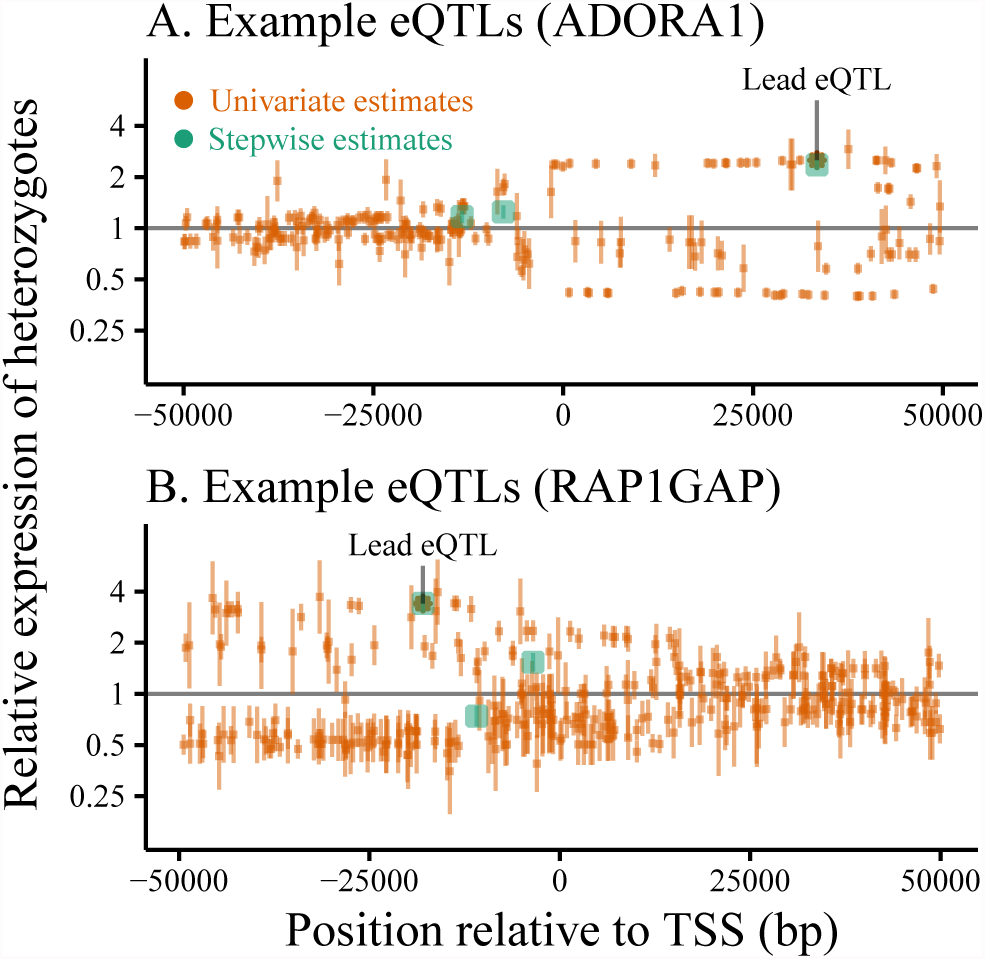
*cis*-eQTLs called using forward-stepwise mapping from whole-blood RNA sequencing in 922 European individuals. eQTL effect sizes are polarized relative to the ancestral allele. Stepwise effect size estimates (green) are compared to those obtained by testing each SNP independently (orange) at two example loci. Vertical bars mark ±2 standard errors around the estimated effect size for each SNP. Lead eQTLs (smallest *p*-value) from independent testing are marked with asterisks.

The median expression of individuals heterozygous for a called, expression-increasing eQTL is 21% higher than that of the expression of homozygotes (-18% for expression-decreasing eQTLs).

When polarizing effect sizes relative to the ancestral allele, we detected 3,284 and 3,303 eQTLs predicted to increase and decrease gene expression, respectively (not significantly different by binomial test; *p* = 0.82). Expression-increasing eQTLs have comparable, though significantly greater effect sizes (in terms of fold-change) than expression-decreasing eQTLs (Fig 4A, *p* = 0.0037 by two-sided Kolmogorov-Smirnov test). In particular, we detected fewer expression-decreasing eQTLs of large effect; there are 139 significant eQTLs predicted to cut gene expression in half, as compared to 243 predicted to double expression.

**Fig. 4.**
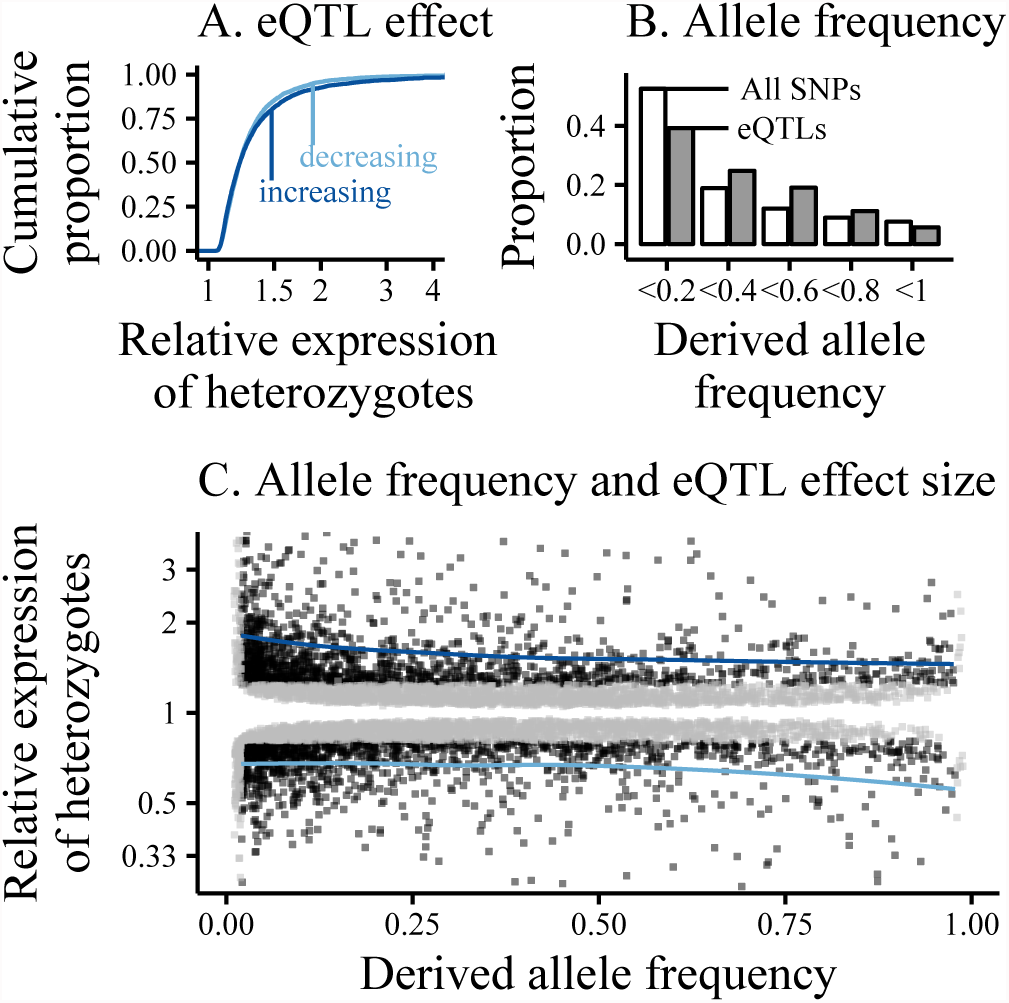
*cis*-eQTLs called using forward-stepwise mapping in 100kb windows centered on the TSS of autosomal, protein-coding genes. Data from whole-blood RNA sequencing in 922 European individuals (DGN). (A) Cumulative distributions of estimated eQTL effect sizes, represented as the ratio of eQTL-heterozygotes to eQTLhomozygotes. Expression increasing eQTLs (derived allele increases expression) are shown in dark blue, expression decreasing eQTLs in light blue. (B) Allele frequency spectrum of eQTLs (grey) relative to the background of all tested SNPs (white) (C) Joint distribution of derived allele frequency and estimated effect size of called eQTLs. Black points show eQTLs with effect sizes that we are powered to detect at all allele frequencies (estimated effect greater than the minimum effect of eQTLs with MAF<0.02), grey points show especially rare eQTLs (MAF<0.02) and eQTLs with effect sizes that we are only powered to detect at higher allele frequencies. Blue lines show the loess-fit of derived allele frequency vs effect size for well-powered expression-increasing and -decreasing eQTLs (black points).

There are both technical and biological explanations for the difference in the effect size distributions of expression-increasing and -decreasing eQTLs. Technically, we may have less power to detect expression-decreasing eQTLs. Biologically, large decreases in gene expression may be less well-tolerated than expression increases.

The joint distribution of allele frequency and effect size shows that, among significant eQTLs, rare SNPs tend to have larger effect sizes than common SNPs (Fig 4C; *p* < 2.2*e* – 16 by Kolmogorov-Smirnov test). Of the 4,949 significant eQTLs with derived allele frequency between 10-90%, the median expression-increasing eQTL is predicted to alter gene expression by 18% (-15% for expression-decreasing eQTLs). However, for the 1,638 significant eQTLs with derived allele frequency less than 10% or greater than 90%, the median expression-increasing eQTL is predicted to alter gene expression by 40% (-25% for expression-decreasing eQTLs). This negative correlation between minor allele frequency and effect size is consistent with the hypothesis that purifying selection acts against *cis*-regulatory variation.

However, these eQTL effect sizes can be difficult to interpret. First, previous work has discussed the challenges inherent in estimating eQTL effect sizes when there are multiple causal variants at a locus (28). Second, power to map eQTLs of the same effect varies with allele frequency. Third, a statistical phenomenon referred to as ‘winner’s curse’ (29, 30) could induce a relationship between allele frequency and estimated effect size. We explore the impacts of each of the above on our eQTL effect size estimates.

First, we observe that univariate (independent) effect size estimates for SNPs are similar to those derived from our stepwise approach (Fig 3). It is therefore unlikely that the presence of multiple causal variants per locus impacts the observed relationship between eQTL effect size and allele frequency.

Variable eQTL-detection power by allele frequency, however, does impact our eQTL calls. One might expect eQTL-based analyses to be limited to relatively common regulatory variants. Indeed, the allele frequency spectrum of our called eQTLs is shifted towards common variants (derived allele frequency between 20-80%) relative to the background distribution of all candidate SNPs (Fig 4B).

Further, as we lack the power to call rare SNPs with small effects as eQTLs, one might expect the median effect size of rare eQTLs to be inflated relative to common eQTLs. However, when considering only eQTLs with large enough effect sizes to be detected at all allele frequencies (details on power thresholds in Supp Table S1), we see that rare eQTLs are still estimated to have larger effects on gene expression than common eQTLs (Table 1). This suggests that decreased power to detect rare, small-effect eQTLs affects the number of eQTLs called at different allele frequencies, but is not sufficient to explain the increased median effect size of rare eQTLs relative to common eQTLs.

**Table 1.** Summary of the effects of variable power and winner’s curse on eQTL effect size estimates and their relationship with minor allele frequency. Effect sizes reported as percent change of eQTL-heterozygotes relative to the homozygous ancestral. To explore the effects of winner’s curse, individuals were split into an eQTL ascertainment set and an effect-size validation set. To explore the impact of variable power to call eQTLs of the same effect across allele frequency bins, eQTLs were filtered to remove those with estimated effects that we would lack power to call at low allele frequencies. Here, we consider eQTLs with estimated effects larger than the minimum magnitude of estimated effect for significant, rare eQTLs (MAF<0.02). To ensure conservative estimates of the relationship between effect size and allele frequency, we also remove rare eQTLs (MAF<0.02) from comparisons across allele frequency bins.

|  | Median eQTL effect |  |  |  |
| --- | --- | --- | --- | --- |
|  | Full data |  | Ascertainment Validation |  |
| expression increasing | 21% |  | 22%<br>18% |  |
| expression decreasing | -18% |  | -18%<br>-14% |  |
|  | MAF 2-10% | MAF >10% | MAF 2-10% | MAF >10% |
| expression increasing filtered for power | 48% | 40% | 52%<br>44% | 45%<br>45% |
| expression decreasing filtered for power | -28% | -28% | -29%<br>-25% | -29%<br>-29% |

However, this relationship between eQTL allele frequency and effect size could also result from winner’s curse. It is expected that, conditional on a variant being called as an eQTL, its effect size will tend to be overestimated (29, 30). This is likely to be particularly true for rare variants.

To test whether the increased effect sizes of rare eQTLs is due to winner’s curse, we ascertained eQTLs and estimated their effect sizes in separate subsamples of the full dataset. In the ascertainment set, eQTLs are estimated to have a median effect of 22% (-18% for expression-decreasing eQTLs) on expression. In the validation set, however, the median eQTL effect is 18% (-14% for expression-decreasing eQTLs; *p* = 9.51*e* –15 by paired t-test; see Supp Fig S1 for comparison of effect sizes estimated in the ascertainment and validation sets).

This systematic decrease in effect sizes estimated in the validation dataset relative to those from the ascertainment dataset demonstrates that winner’s curse inflates effect size estimates during eQTL ascertainment. Comparison of median effect size estimates in the ascertainment and validation sets for more rare (0.02<MAF<0.1) and more common (MAF>0.1) eQTLs, respectively, reveals that winner’s curse disproportionately inflates effect size estimates at the low end of the allele frequency spectrum (Table 1).

Indeed, effect size estimates obtained in the validation set no longer show any relationship between eQTL allele frequency and effect size (Supp Fig S1B). This is consistent with Tung et al. 23, who also see a significant impact of winner’s curse on the effect size estimates of rare eQTLs. We therefore conclude that eQTL effect sizes alone fail to provide evidence for constraint on gene expression.

Next, we wanted to understand whether our analyses of eQTL effect sizes provided evidence of the absence of constraint, or merely the absence of evidence. We therefore used allele-specific expression (ASE) to validate the effect size estimates of called eQTLs and test the relationship between eQTL effect size and minor allele frequency in an independent dataset.

### eQTL effects on allele-specifc expression

Allele-specific expression (ASE) analysis relies on a heterozygous site in a transcript to identify reads derived from each allele within an individual. These allelic read counts are then used to quantify expression levels from each haplotype.

Variation in expression across haplotypes captures the sum of the regulatory effects of all *cis*-heterozygous sites in that individual, regardless of their frequencies in the population. Therefore, ASE-based analyses incorporate the effects of genetic variants that are too rare to be detected in eQTL studies. On the other hand, eQTL-mapping is better suited to estimate the effect of each individual variant on gene regulation. ASE and eQTL mapping are therefore complementary tools for measuring *cis*-gene regulatory genetic variation.

To expand on our previous findings and to sidestep the statistical challenges inherent in eQTL effect size estimation, we combined our called eQTLs with allele-specific expression (data from GTEx version 6, GTEx Consortium 21). We measured allelic imbalance using a squared Z-score of the number of reads containing the alternative allele in 372,473 geneindividual-tissue trios (e.g. a gene expressed in a given tissue in a given individual) that passed our QC filters (see). To determine whether our estimated eQTL-effects are reflected in ASE, we analyzed allelic imbalance at genes for which we called an eQTL.

On average, in genes with at least one significant eQTL, individuals heterozygous for an eQTL have higher allelic imbalance than individuals who are homozygous for all called eQTLs (Fig 5A). In addition, the effect sizes of individual eQTLs are correlated with the allelic imbalance of individuals that are heterozygous for those eQTLs (Fig 5B). This echoes previous work demonstrating that ASE measurements are generally consistent with mapped *cis*-eQTLs at a locus (31).

**Fig. 5.**
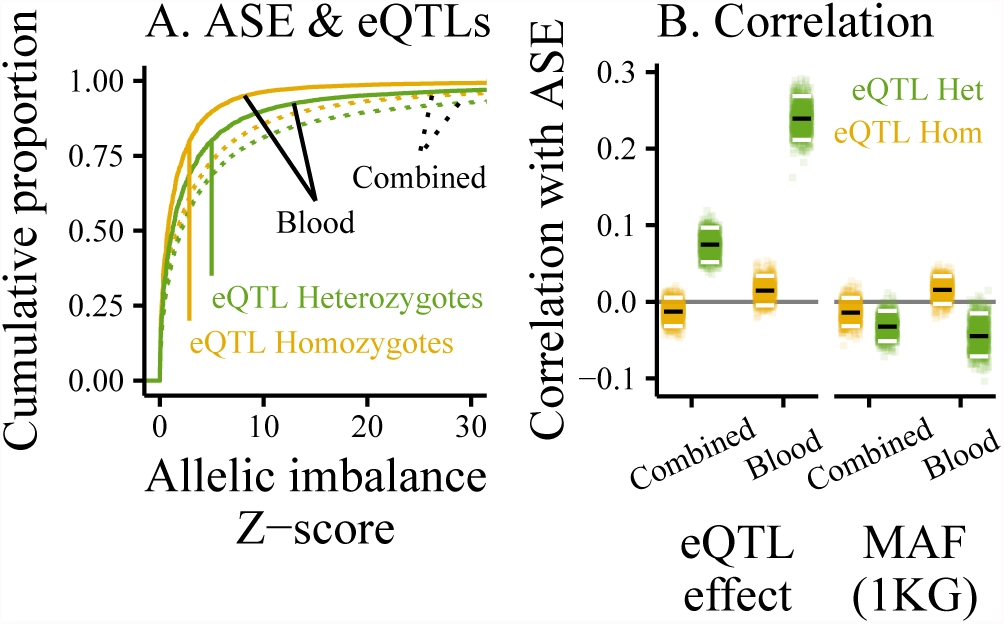
eQTL effects reflected in allelic imbalance of genes with at least one signifcant eQTL. ‘Blood’ shows ASE in whole blood, ‘Combined’ shows combined-tissue ASE, measured by summing reads spanning each ASE site in the same individual across all tissues sampled. (A) Cumulative distributions of ASE measured in whole blood (solid) and combined-tissue ASE (dotted), separated into eQTL-heterozygotes (green), and -homozygotes (homozygous for all called eQTLs, yellow). (B) Rank correlation between allelic imbalance and eQTL effect (left) or minor allele frequency in 1000 Genomes (right). The effect/MAF of the most significant heterozygous eQTL (or, for eQTL-homozygotes, the most significant eQTL) was used. To determine the robustness of these correlations, each point shows the rank correlation in one of 1000 samples generated by bootstrapping over genes. Median correlations of bootstrap samples marked in black, 95% quantiles in white.

To gain additional insight into the structure of the gene regulatory landscape across tissues, we compared a single-tissue analysis of blood to a combined-tissue analysis in which all reads spanning an ASE site in an individual were summed across all tissues sampled.

Interestingly, regardless of eQTL-status (-homozygote or -heterozygote), the distributions of combined-tissue ASE Z-scores are right-shifted (larger) than ASE Z-scores measured in blood (Fig 5A). This shift can be explained by differences in read depth; increased read depth decreases sampling variance, thus increasing our ability to detect bona fide allelic imbalance (and increasing the resulting ASE Z-score). Due to the addition of reads sampled across tissues, the average read depth per ASE site is quite different between the combined-tissue and blood-specific data (median 133 reads for combined-tissue and 28 reads for blood, more detailed read depth information in Supp Fig S6A). Therefore, increased confidence to detect ASE likely drives the bulk of the larger Z-scores seen in the combined-tissue analysis.

It is also worth noting that the combined data will have increased Z-scores only if allelic imbalance is concordant across tissues. Like many other eQTL and ASE analyses (e.g., GTEx Consortium 21, Wheeler et al. 32, Flutre et al. 33), our findings suggest that many gene regulatory effects are consistent (or, at least, are unlikely to be inconsistent) across tissues.

Finally, we explored whether the allelic imbalance data reveal evidence of selection acting against large-effect eQTLs (Fig 5B). In both combined-tissue and blood-specific ASE, there is a subtle, but significant negative correlation between the allelic imbalance of individuals that are heterozygous for a called eQTL and the allele frequency of that eQTL. This subtle negative relationship between allele frequency and effect size suggests that selection on eQTLs is present, but weak.

### Effects of *cis*-regulatory variants vary with allele frequency

We next examined signals of constraint on variants that are too rare to be detected in eQTL studies and on variants in *cis* to genes without significant eQTLs. To do this, we first sought to relate observed allelic imbalance to the amount of *cis*-regulatory variation at a locus. In our model, each *cis*-heterozygous site contributes to the expected variance in the ratio of reads expressed from each allele (detailed in). In this case, we would expect the number of *cis*-heterozygous sites to be linearly related to our ASE Z-score.

We therefore estimated the contribution of an average *cis*-heterozygous site to ASE using linear regression across all individual-gene pairs. In individual tissues and in the combined-tissue analysis, we observed significantly positive relationships between number of *cis*-heterozygous sites and allelic imbalance at a locus (Fig 6). This relationship between genetic variation and allelic imbalance suggests that variance in gene expression has a genetic basis, rather than being solely driven by sampling noise or environmental factors.

**Fig. 6.**
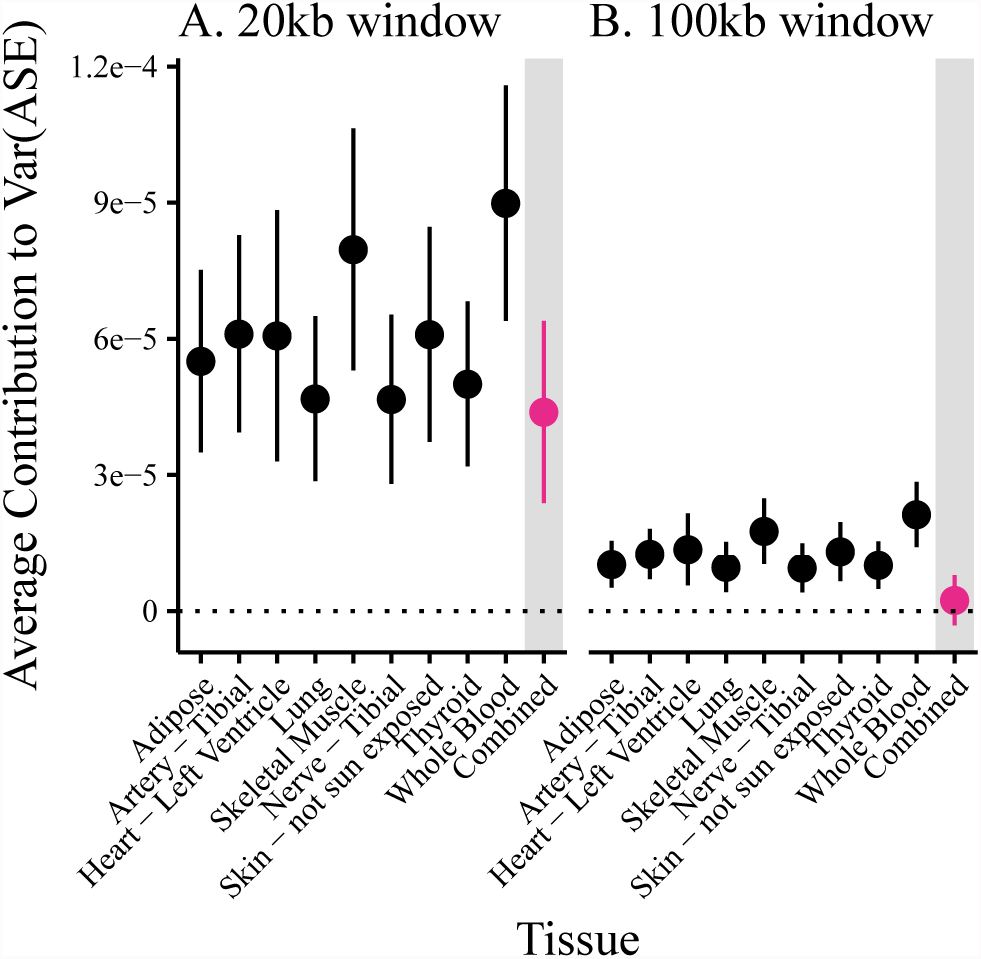
Average effect of a *cis*-genetic variant on allelic imbalance. Estimates are based on linear regression of an individual’s number of *cis*-heterozygous sites within (A) 10kb and (B) 50kb of the TSS of a gene and measured ASE for each of the nine best sampled GTEx tissues. Vertical lines show 95% confidence intervals, estimated using a weighted jackknife as described in (34). For reference, the dotted line marks zero estimated effect.

**Fig. 7.**
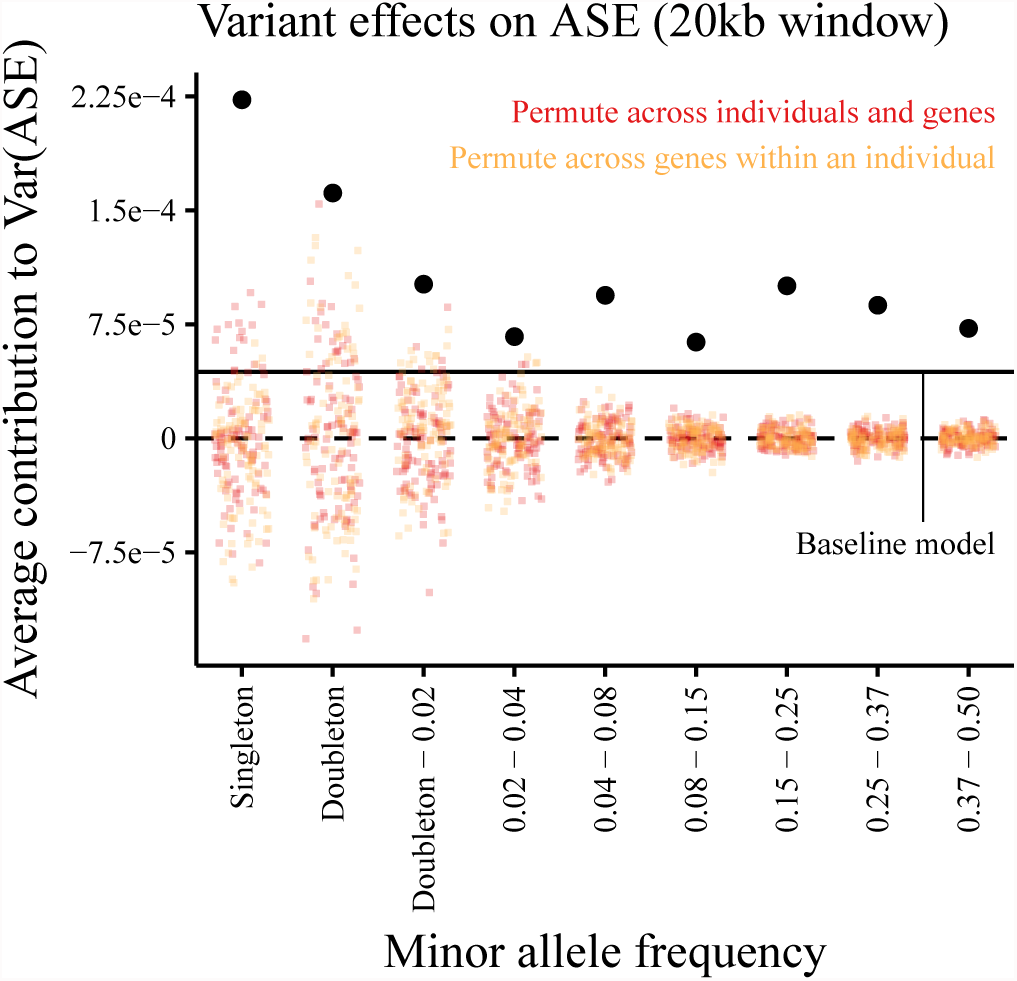
Effects of *cis*-genetic variants on allelic imbalance given their allele frequencies. Estimates are obtained from multiple regression on allelic imbalance using the number of *cis*-heterozygous sites in each allele frequency bin as predictors. Here, we include sites in a 20kb window centered on the transcription start site of a gene with measured allelic imbalance. Estimates from combined-tissue ASE (reads spanning each ASE site in a single individual are summed across all tissues sampled) are shown in black. Colored points represent estimates from 100 permutations of (1) ASE measurements across genes and individuals (red) and (2) ASE measurements across genes within an individual (orange). Horizontal lines mark zero effect (dashed) and the average variant effect estimated by regressing allelic imbalance on the total number of *cis*-heterozygous sites (solid).

Interestingly, the estimated per-variant effect on ASE strongly depends on the size of the region that we consider to be in *cis*. In a 20kb window, per-variant effect size estimates are similar across tissues and are all significantly positive. However, in a 100kb window, the estimated per-variant effect in the combined-tissue analysis is indistinguishable from zero (Fig 6).

This likely reflects the spatial distribution of causal regulatory variants. Similar to what has been found in prior work (35, 36), we find that eQTLs are enriched near the transcription start sites of genes (Supp Fig S2). If this is also true for rarer gene-regulatory variants, increasing the size of the genomic region will decrease the proportion of causal variants in the model and so decrease the average effect size per variant.

Next, we expanded our model to incorporate allele frequency information. If purifying selection acts to keep large-effect regulatory variants rare, we would expect that, on average, rare *cis*-heterozygous variants should contribute more to allelic imbalance than common *cis*-heterozygous sites. For each individual-gene pair, we counted the number of heterozygous *cis*-regulatory sites in each of ten allele frequency bins. We then used multiple regression to estimate the average contribution of *cis*-heterozygous sites in each allele frequency bin to observed allelic imbalance. We find that the average contribution of a singleton in this dataset to the variance in ASE is approximately 2.5× greater than that of a very common variant.

Permutation of ASE measurements across individuals and genes, as well as across genes within an individual, suggest that there is a significant relationship between the allele frequencies of genetic variants and allelic imbalance.

Qualitatively, the relationship between allele frequency and effect on allelic imbalance persists when considering genetic variants in a much larger candidate *cis*-regulatory region (100kb window shown in Supp Fig S4). As in the case of the average per-variant effect (Fig 6), increasing the size of the *cis*-regulatory region decreases the estimated effect for variants in each allele frequency bin. However, the contributions of rare variants (singletons and doubletons) remain greater than those of common variants.

Importantly, the relationship between allele frequency and contribution to allelic imbalance is so subtle that it is detectable only when reads are combined across tissues; single-tissue analyses show no clear relationship between allele frequency and estimated effect size (examples whole blood and skeletal muscle shown in Supp Fig S5). This is likely due to the fewer genes, fewer individuals sampled per gene, and lower read depth per individual-gene pair in single tissues as compared to the combined set (for sample size comparisons, see Supp Fig S6; for read depth comparisons, see Supp Fig S7).

From this, we conclude that constraint on gene expression does reshape patterns of regulatory genetic variation. However, the subtlety of this signal suggests that selection on gene expression is relatively weak.

### Proportion of heritability explained by rare variants

Another way to understand the strength of selection on gene expression is to ask what proportion of the heritability (or, the proportion of the genetic variance 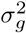) of gene expression can be explained by rare variants.

Two components determine how much genetic variance is explained by a variant: the variant’s squared effect size and its allele frequency (particularly its contribution to heterozygosity, 2*pq*). When considering the total proportion of heritability explained by all variants at a given allele frequency, the allele frequency spectrum (the proportion of variants in each allele frequency bin), is a third key factor.

Many polygenic trait models assume that all variants, regardless of their allele frequency, contribute equally to genetic variance (2*pqß*^2^; e.g., Yang et al. 37, Bulik-Sullivan et al. 38). As their low allele frequencies cause rare variants to contribute less to heterozygosity (2*pq*), this type of model implicitly assumes that rare variants have larger effects (*β*^2^) than common ones.

Others (e.g., Speed et al. 39) more explicitly account for the relationship between allele frequency (and corresponding differences in LD) and variance explained. For a more detailed comparison of heritability estimation and variance partitioning methods, see Evans et al. 40.

While these models were not designed with singletons in mind, their varying assumptions about allele frequency, effect size, and heritability led us to consider what effect the observed 2.5× relative effect size difference between rare and common variants would have on the proportion of heritability explained by rare variants.

To explore this, we combined our effect size estimates with allele frequency spectra at each gene to estimate the total variance in allelic imbalance explained by our ASE-model of genetic variance (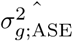). We then used the same effect size estimates and allele frequency spectra to estimate the proportion of 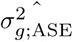 explained by each allele frequency bin (summary of allele frequency spectra in Supp Fig S8). Across genes, we find the median heritability explained by all singletons (in a sample of 122 individuals) to be a mere 5% (Fig 8). Table 2 shows comparisons of our model to two extreme cases: (1) rare and common variants contribute equally to genetic variance (2*pqß*^2^), (2) rare and common variants have equal effects on gene expression (*ß*^2^).

**Fig. 8.**
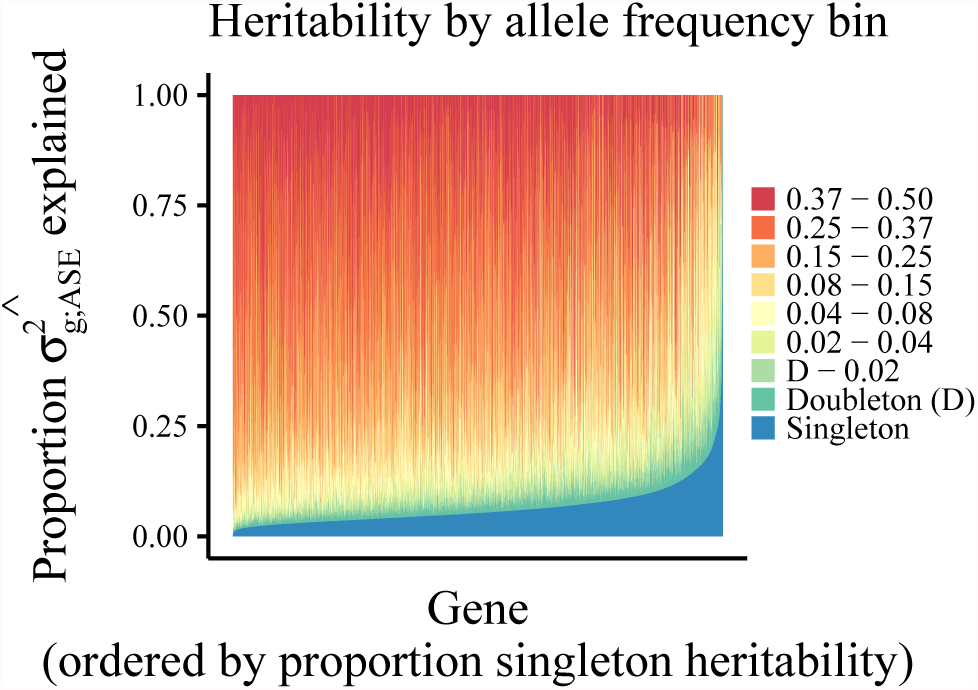
Partitioning the heritability of allelic imbalance by allele frequency. For each gene, we combine the average heterozygosity (2*pq*) and number of segregating sites in each allele frequency bin with the estimated average effect size of variants in that bin to calculate the total heritability of allelic imbalance captured by our ASE-model (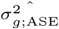). We then calculate the proportion of 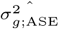 explained by each allele frequency bin. Each vertical bar represents a single gene for which we measured ASE, colors represent the proportion of that heritability attributed to variants in each allele frequency bin.

**Table 2.**
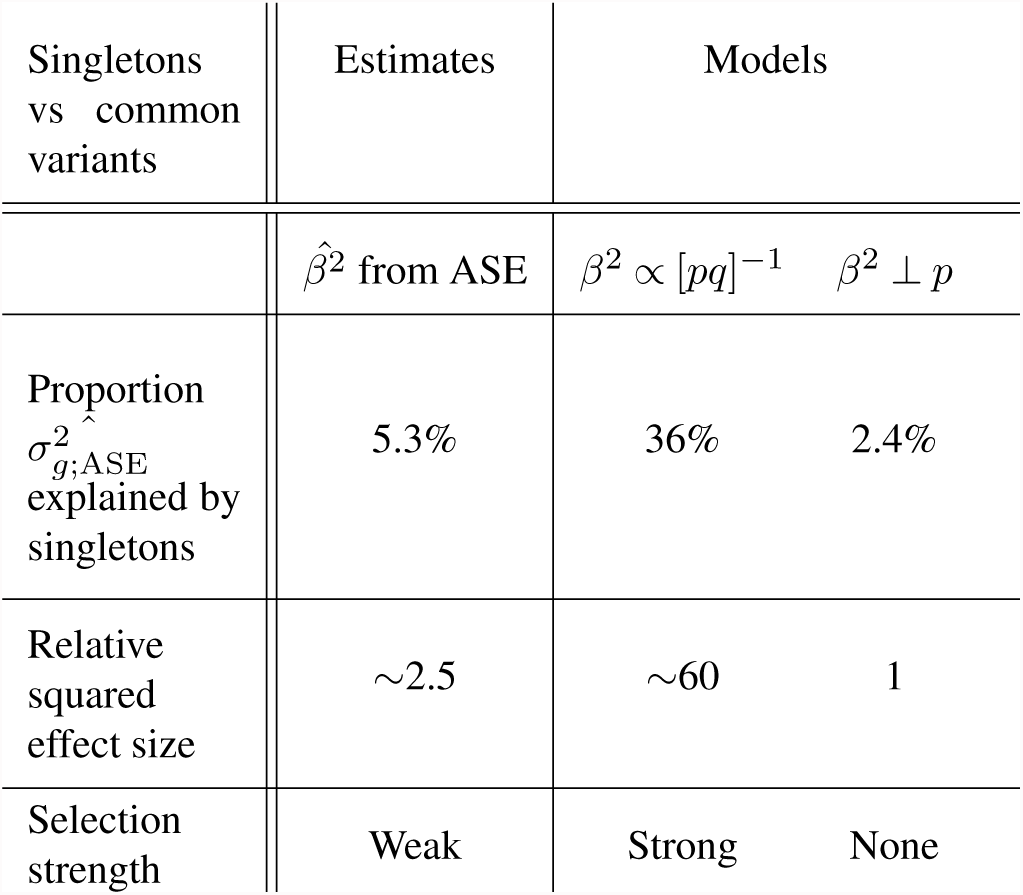
Singleton contributions to heritability based on GTExV6 ASE data (*cis*-regulatory region ±10kb from TSS). ‘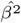 from ASE‘ shows estimates of the median proportion of 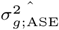 explained by singletons based on effect sizes estimated from our ASE-model. ‘*β*^2^ ∝ [*pq*]^−1^‘ shows the median expected per-gene heritability explained by singletons if singletons and common variants contributed equally to genetic variance. This is driven by the fact that 35% of all variants (median 36% of variants per gene) are singletons. The large relative effect size required for singletons to explain this proportion of heritability reflects the low variance contribution of rare variants as a consequence of their low allele frequency (2*pq*). ‘*ß*^2^ ⊥ *p*‘ shows estimates of the median proportion of heritability explained by singletons if effect size and allele frequency were independent (i.e. singletons and common variants had equal effects on gene expression).

As a result of their increased effect sizes relative to common variants, our model suggests that singletons explain more of the heritability of gene expression than expected under neutrality (effect size independent of allele frequency). However, in these data, the relative effect size difference between singletons and common variants (0.37<MAF<0.5) is far less than what would be required for a singleton to contribute as much as a common variant to heritability.

It is important to note that changing sample size may alter the proportion of heritability explained by singletons in several ways. Singletons in larger samples will be more rare in the population than singletons in a smaller sample. Owing to the recent, rapid growth of the human population, singletons in larger samples will also be more abundant in terms of the proportion of total variants (41–43).

The increased abundance of singletons in larger sample sizes might suggest that the total proportion of variance explained by singletons should increase. However, each of these new, lower-frequency singletons will have correspondingly lower heterozygosity (2*pq*) and lower heritability contributions as a result. Finally, the relative effect size between singletons and common variants will likely also change with changing allele frequency. The complex interaction between these factors makes it difficult to directly compare estimates of the heritability explained by singletons across datasets (see Hernandez et al. 25 for another approach).

Interestingly, although the median heritability explained by singletons is relatively small, there is significant variation in singleton-heritability across genes (Fig 8). As our singleton effect size estimate is constant across genes, this variation is due to differences in allele frequency spectra at different genes.

This variation led us to wonder how much of the constraint on gene expression (i.e. relative effect size differences between rare and common variants) that we observe is driven by a subset of particularly tightly constrained genes.

### Patterns of eQTLs and ASE suggest variable constraint on expression across gene classes

To start to untangle variation in the strength of constraint on gene expression across the genome, we compared patterns of regulatory variation across gene sets.

We expect genes with greater constraint on expression to have lower levels of *cis*-regulatory variation than genes with less-constrained expression. To test whether this is the case, we compared patterns of eQTLs across gene sets predicted to have varying tolerance to large changes in gene expression. Specifically, we utilized Probability of Loss-of-Function Intolerance (*pLI*) (13). *pLI* measures, for each gene, the relative depletion of protein-truncating variants (PTVs) observed in healthy individuals compared to the expectation under a detailed mutation model.

Protein-truncating variants are expected to decrease gene expression by half. Certain genes may therefore be depleted for PTVs because they are intolerant to such large expression changes. We stratified genes into two classes based on their *pLI*-predicted level of constraint on expression (low: *pLI*<0.1, high: *pLI*>0.9) and tested for differences in *cis*-regulatory variation.

In total, 4,401 eQTLs were mapped around 3,103 unique low-*pLI* genes (of 6,813 genes tested) and 688 eQTLs were mapped around 511 high-*pLI* genes (of 2,576 genes tested). We detected fewer eQTLs per gene for dosage sensitive, high*pLI* genes than for less constrained, low-*pLI* genes (Fig 9A). These differences in the number of *cis*-regulatory sites led to significant differences in the amount of genetic variance (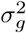) from eQTLs for low- and high-*pLI* genes (Fig 9B; *p* < 2.2*e* –16 by two-sided Kolmogorov-Smirnov test). For each individual in our dataset, we estimated the genetically regulated component of expression based on their genotypes at called eQTLs. We then measured genetic variance by taking the standard deviation of predicted expression across individuals.

**Fig. 9.**
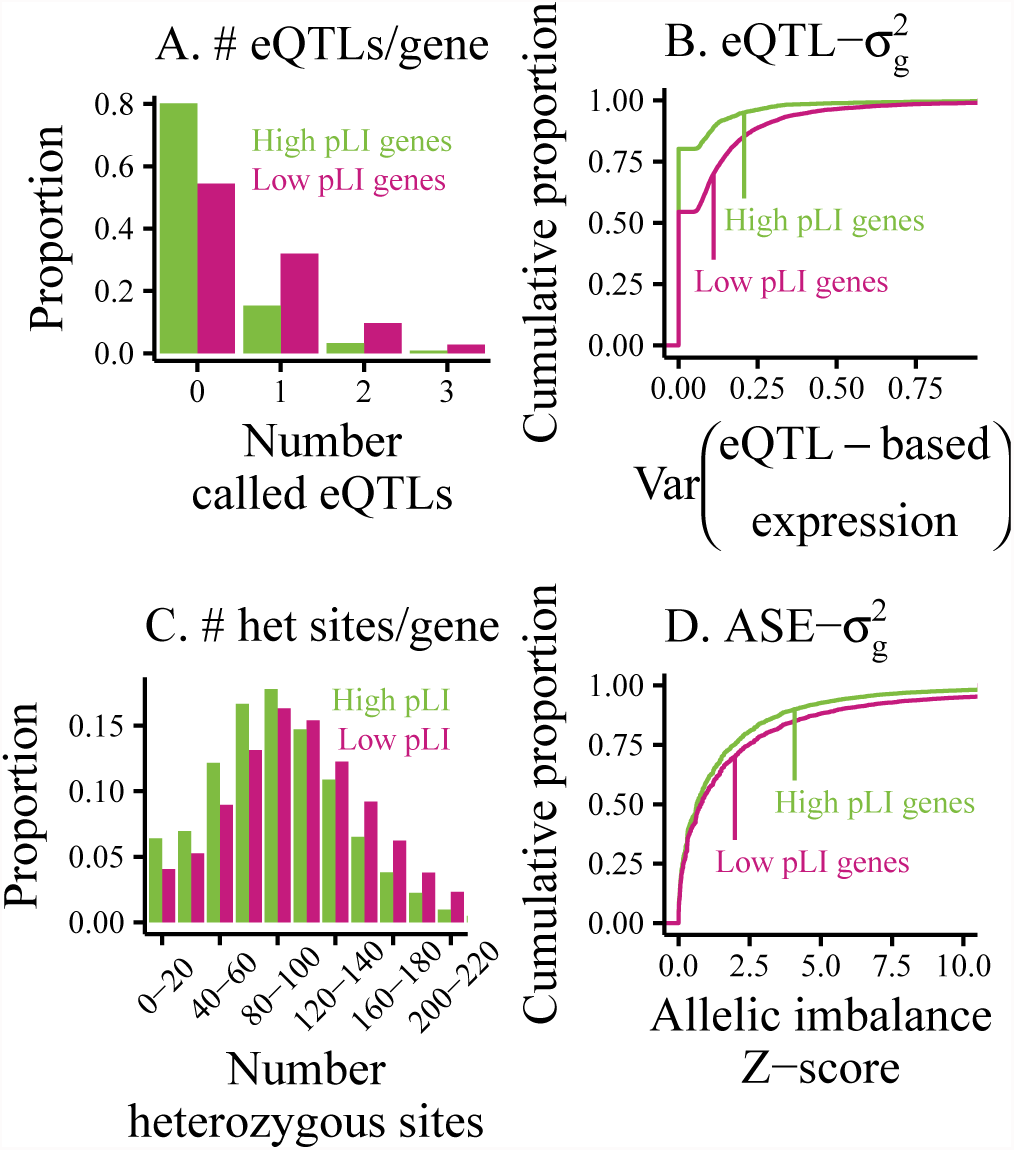
Regulatory genetic variation across gene sets with varying levels of constraint on gene expression as predicted by *pLI* class. (A) Histogram of the number of *cis*-eQTLs called per gene by *pLI* class. (B) Cumulative distributions of eQTL-estimated genetic variance by *pLI* class. For each gene in each individual, expression levels were predicted using genotypes and effect size estimates of called eQTLs. Genetic variance was then estimated as the standard deviation of eQTL-based expression levels across individuals. (C) Histogram of the number of *cis-*heterozygous sites within 50kb of the TSS by *pLI* class. (D) Cumulative distributions of allelic imbalance, measured in whole blood, by *pLI* class.

To quantify differences in genetic variance across genes, we compared the median eQTL-estimated genetic variance for genes in each *pLI* class. We then assumed log-normally distributed expression values to estimate ranges of gene expression in the population.

Across genes with at least one called eQTL, we estimate that 95% of individuals possess genetic variation resulting in expression levels between 0.83 – 1.21 × the population mean. However, for the median low-*pLI* gene, these ranges are 0.82 –1.22 × and, for the median high-*pLI* gene, they are 0.85 – 1.18×. This may suggest that constraint on expression levels removes more regulatory genetic variation around dosage-sensitive high-*pLI* genes than less sensitive low-*pLI* genes.

Importantly, this observation could also be explained by differences in background selection across gene sets. In particular, high-*pLI* genes experience greater levels of coding sequence constraint than do low-*pLI* genes. We might therefore expect background selection to decrease the amount of available regulatory genetic variation around high-*pLI* genes.

As shown in Supp Fig S2A, there are indeed fewer common SNPs in *cis* to low-*pLI* genes (MAF>0.05, within 50kb of the transcription start site) than to high-*pLI* genes. SNPs around low-*pLI* genes also tend to be slightly more common than those around high-*pLI* genes (Supp Fig S2C), and low-*pLI* eQTLs tend to be closer to the TSS (Supp Fig S2B/D). While these trends point to a role for background selection in shaping patterns of genetic variation around genes, they are unlikely to explain the striking differences observed in called eQTLs.

Differences in genetic variance across genes can also be seen in allele-specific expression. As in the comparison of ASE across tissue types, our ASE Z-score increases with increasing read depth. As high-*pLI* genes tend to be more highly expressed, and therefore have more sampled reads than low*pLI* genes (read depth information in Supp Fig S7), we might expect to more confidently call ASE (i.e. higher Z-score) in high-*pLI* genes. However, we observed the opposite pattern; ASE Z-scores for high-*pLI* genes are left-shifted (lower) relative to those for low-*pLI* genes (Fig 9D).

In blood, 20.9% of 74,262 individual-gene pairs involving low-*pLI* genes have a Z-score of allelic imbalance greater than three. This level of allelic imbalance is unlikely to arise from read sampling, suggesting that each haplotype is subject to different *cis*-regulatory effects. By contrast, only 15.6% of 36,715 individual-gene pairs involving high-*pLI* genes have Z-scores greater than three (significantly different by binomial test, *p* < 2.2*e*16).

Although the differences in read depth suggest that the downward shift in the distribution of high-*pLI* ASE Z-scores relative to low-*pLI* scores under represents the difference in gene expression variance between these gene classes, these patterns of allelic imbalance point to increased expression constraint on dosage-sensitive, high-*pLI* genes.

The lower gene expression variance seen in high-*pLI* ASE is reflected in patterns of genetic variation; high-*pLI* genes tend to have fewer *cis*-heterozygous sites than do low-*pLI* genes (Fig 9C). These findings suggest a link between expression variance and genetic variation. Combined with the differences in genetic variance across gene sets, this could be interpreted as evidence of selection reducing the variance in the expression of tightly constrained genes by reducing the number of segregating *cis*-regulatory sites. In this case, however, these differences in genetic variation across gene classes are difficult to disentangle from the effects of background selection. These stark differences in regulatory variation across gene sets suggest that constraint in gene expression varies widely across the genome. The marked variation in the proportion of genetic variance explained by singletons (Fig 8) also suggests the presence of tightly constrained genes.

However, it remains unclear how much of the genome-wide signal of constraint on regulatory variants can be attributed to strong constraint on a subset of genes, like high-*pLI* genes, and how much is due to weak constraint on expression subtly shifting the joint distribution of allele frequency and regulatory effect genome-wide. The extremely small number of genes with large contributions from rare variants (87% of genes have <10% genetic variance explained by singletons, 98% have <20%) may suggest that the bulk of the relationship between allele frequency and effect size is driven by weak selection on many genes, but further work is required to corroborate this interpretation. In particular, larger datasets of paired whole genome and RNA sequencing will increase the sampled genetic diversity between individuals at a locus and provide additional insight into the landscape of selection on gene expression.

## Discussion

Our analyses of eQTLs and ASE suggest that selection on gene expression has limited effects on gene regulatory variants. However, analyses of loss-of-function and copy number variants suggest that selection on gene expression changes of 1.5× is, at least in some cases, sufficient to alter patterns of gene expression variation.

These observations could be reconciled by arguing that the 1.2× expression changes caused by the average eQTL and the 1.5× changes in expression caused by a gene duplication have very different effects on fitness. One might also argue that the effects seen in loss-of-function and copy number variant analyses are driven by a small set of tightly constrained genes, or that loss-of-function and copy number variants have effects on fitness that are unrelated to their effects on gene expression. These arguments might lead us to conclude that gene expression has limited downstream effects on human traits and on fitness. However, even if gene expression affects fitness, if gene regulation is highly polygenic and/or pleiotropic, selection may be unable to cause dramatic changes in the frequencies of variants that affect gene expression.

Rampant polygenicity is likely to decrease the effectiveness of selection on individual regulatory variants because if, at a single locus, many variants can affect gene expression, a new mutation with a large effect may arise on a background with existing, compensatory genetic variation. This would reduce the effective selection coefficient on the new mutation. Such background-effects may explain how the expression of common copy number variants is similar between individuals with two and three gene copies.

Additionally, if gene regulation is highly polygenic and spatially structured, a significant eQTL may not always reflect a single causal variant with the effect size estimated by eQTL mapping. Instead, it may represent a composite of several causal sites, each of which has a small effect on expression. If multiple causal variants with concordant effects happen to be in linkage disequilibrium in a sample, their effects may be aggregated into a single “synthetic” eQTL with a large effect (for more on synthetic associations, see Dickson et al. 44). If the variants that comprise such a synthetic eQTL are not in linkage disequilibrium in the population, their effect sizes may individually be of sufficiently small effect to be nearly neutral, even if selection would be strong enough to act on the magnitude of change attributed to the group. Future work is required to determine to what degree eQTLs may represent groups of concordant, small-effect variants.

Further, this challenge of polygenicity is not limited to the regulation of a single gene. If a fitness-relevant trait is affected by the expression levels of many genes, the fitness consequence of a regulatory variant that affects the expression of a single gene is likely to be small.

Pleiotropic effects of regulatory variants may also decrease the effectiveness of selection. For example, if a single variant affects the expression of multiple genes, and the resulting expression changes each have an effect on fitness, the selection pressure experienced by the variant will be a combination of the constraint on each expression trait with which it is associated.

Such regulatory pleiotropy may also arise if the expression of a single gene affects multiple fitness-relevant traits. This is particularly true when considering that regulatory variants and altered gene expression levels may have different effects in different tissues. A variant that affects the expression of a single gene, then, may still have multiple (and possibly opposing) effects on fitness. Interestingly, as the polygenicity of fitness-relevant traits (e.g., the number of genes whose expression modulates the trait) increases, the likelihood that a single regulatory variant causes pleiotropic effects on multiple traits also increases.

In summary, a complex, pleiotropic gene expression network may make it difficult for selection to precisely alter the allele frequency of a single regulatory variant, even when its effects might be large and deleterious from the perspective of one gene expression trait in one tissue context. While there is much work to be done to connect gene expression levels to the one, or many, phenotypic traits seen by selection, we conclude that constraint has subtle, but detectable, effects on the genetic architecture of gene regulation.

## Materials and Methods

### Expression of gene duplicates

We identified loci containing copy number variants (CNVs) in healthy individuals, as well as the number of gene copies per locus per individual, by applying LUMPY (45) and Genome STRiP (46) to whole blood RNA sequencing data from version 4 of the Genotype Tissue Expression Project (GTEx) (21). Only CNVs containing an entire protein coding sequence were retained for downstream analysis. We obtained gene expression (RPKM) measurements for each CNV in each individual across 12 tissues.

### eQTL mapping

We obtained genotype and RNA sequencing data from 922 European individuals included in the Depression Genes and Networks (DGN) dataset (20, 47). Genotypes were imputed to 1000 Genomes as described in Kukurba et al. 48. We then polarized genotypes relative to the human ancestral allele.

To determine the human ancestral allele, we compared the human major allele to the human-chimp ancestral allele. We obtained human major alleles from the 1000 Genomes dataset (49) and inferred the human-chimp ancestral allele from Ensembl multiple alignments using Ortheus (50).

At SNPs for which the human major allele and the human-chimp ancestral allele agree, we defined the human ancestral allele as the agreeing allele. At SNPs for which the human major allele and human-chimp ancestral allele disagree and the human minor allele is rare (<5% frequency), we defined the human ancestral allele as the human major allele. At SNPs for which the human major allele and human-chimp ancestral allele disagree and the human minor allele is common (>5% frequency), we defined the human ancestral allele as the human-chimp ancestral allele. At SNPs which lacked data regarding the human-chimp ancestral allele, we defined the human ancestral allele as the human major allele.

We considered SNPs with minor allele frequency greater than 1% within a 100kb window centered on the most upstream annotated transcription start site to be candidate *cis*-eQTLs for the corresponding gene.

To obtain comparable gene expression measurements across individuals, we normalized read counts at each gene by the total reads sampled per individual and log_2_ transformed the resulting measurement.

To allow multiple independent eQTLs per gene, we mapped eQTLs using forward stepwise regression. For a detailed treatment of the challenges introduced by multiple regulatory variants per locus, see (28).

If any SNPs were significantly associated with gene expression (*α* = 0.05 after Bonferroni correction for the original number of candidate eQTLs for the gene), the most significantly associated SNP was added to the model. All SNPs in linkage (*r*^2^ > 0.8) with the newly called eQTL were excluded from the list of candidate eQTLs. This model selection procedure was repeated until no significantly associated SNPs remained.

### Modeling Allele-Specific Expression (ASE)

In general, ASE is measured by comparing reads expressed from each allele at a particular locus within an individual. This relies on a heterozygous site within a transcript that can be used to identify the allelic origin of each read.

Here, we modeled allele specific expression of the *g*^th^ locus in the *i*^th^ individual as a squared Z-score of reads containing the alternative allele 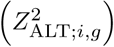. We used the squared score because it is agnostic to the direction of allelic bias (excess of reference or of alternative alleles).

Under a null model in which read counts follow a Binomial distribution,
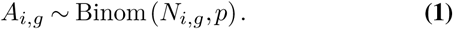

*A_i,g_* is the number of sampled reads containing the alternative allele, *p* is the underlying proportion of reads that contain the alternative allele, and *N_i,g_* is the total number of reads sampled at the locus. The expected number of sampled reads containing the alternative allele is then,
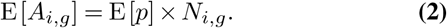

In the absence of locus- and allele-specific *cis*-regulation, the expected proportion of reads expressed from the alternative allele is 0.5. However, due to reference mapping bias, the observed proportion of alternative reads will be slightly lower. This ratio (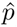) was estimated empirically using global reference bias.
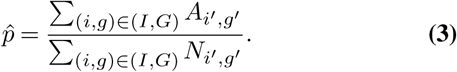

where (*I, G*) is the set of individual-gene pairs at which we measured ASE. The variance in alternative allele counts is therefore,
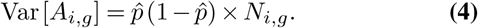

Our ASE statistic is therefore,
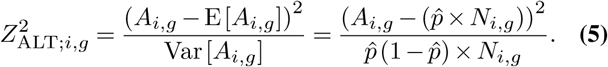

### Contribution of cis-regulatory variants to ASE

Thus far, we have assumed that the underlying proportion of alternative-allele reads at all loci is determined by reference bias alone. To explore the relationship between ASE and locus-specific regulatory variation, we must allow the true proportion of alternative-allele reads to vary across ASE sites. We therefore extended our model with the assumption that, within an individual, any heterozygous site in *cis* to an ASE site can alter expression levels across haplotypes and contribute to allelic imbalance.

Specifically, in each individual *i* at each locus *j*, we assumed the underlying proportion of reads containing the alternative allele (*p_i,g_*) to be Beta distributed. The mean proportion of alternative allele reads is determined by reference bias, as described above. However, we modeled the variance in the proportion of reads expressed from the alternative allele as a function of the number of heterozygous sites in *cis* to each measured ASE site in each individual.
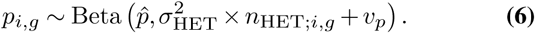

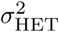 is the variance in the proportion of alternative allele-reads contributed by a *cis*-heterozygous site, *n*_Het;*i,g*_ is the number of *cis*-heterozygous sites at the *g*^th^ locus in the *i*^th^ individual, and *v_p_* is the variance in the proportion of reads expressed from each allele contributed by factors other than *cis*-genetic variation. This could represent expression variance from *trans*-acting or environmental factors.

Note that 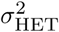 is a genome-wide parameter; this assumes that, across loci, variable sites have similar effects on allelic imbalance. This model also assumes that each *cis*-heterozygous site contributes additively to the variance in the underlying proportion of reads expressed from the alternative allele (Var[*p_i,g_*]).

We then assumed that the number of alternative allele-reads sampled in individual *i* at locus *g* (*A_i,g_*) results from binomial sampling around the proportion of alternative allele reads at that locus.
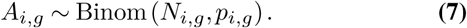

In total, the observed number of reads containing the alternative allele will be Beta-Binomially distributed. The variance in alternative-allele counts is then,
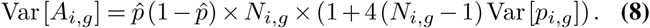

Under this model, we can update our squared Z-score of alleleic imbalance (Eq 5) to account for variance contributed by *cis*-regulatory genetic variation.
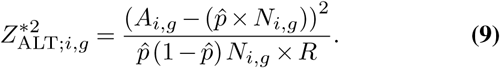

where *R* is
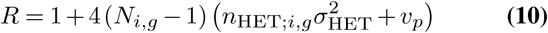

Note that
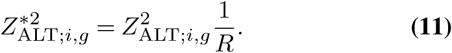

When the number of sampled reads at a locus, *N_i,g_*, is large, 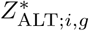, is standard-normally distributed. Regardless of the number of sampled reads, 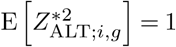. Therefore,
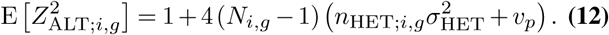

Rearranging Eq 12, 
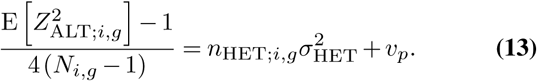

As 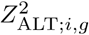, *N_i,g_*, and *n*_Het;*i,g*_ are measurable in data, we can apply linear regression according to the above model to estimate 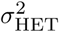.

#### Variation in cis-regulation by allele frequency

The above model assumes that all *cis*-heterozygous sites contribute equally to allelic imbalance. However, if stabilizing selection acts on gene expression, we would expect rare variants to contribute disproportionately to allelic imbalance. We therefore extended our model to allow different values of 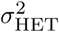 for *cis*-heterozygous sites with different allele frequencies. Now,
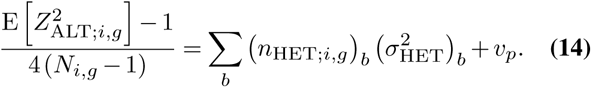

where *b* indexes allele frequency bin. (*n*_Het;*i,g*_) *_b_* is, for individual *i* at locus *g*, the number of *cis*-heterozygous sites with a population frequency that falls in allele frequency bin *b*, and 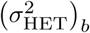 is the variance in allelic imbalance explained HET)by each variant in allele frequency bin *b*.

We can then estimate values of 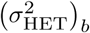 jointly using multiple regression with the counts of *cis*-heterozygous sites in each allele frequency bin as predictors.

### Data analysis of ASE and the effects of *cis*-regulatory variants

In this study, we analyzed ASE in 122 self-reported European individuals with RNA sequencing data and genotype calls from whole-genome sequencing from the Genotype Tissue Expression Project (21). We included the nine best-sampled tissues (whole blood, subcutaneous adipose, tibial artery, heart - left ventricle, lung, skin - not sun exposed, tibial nerve, skeletal muscle, and thyroid) in our analyses.

For an allelic imbalance measurement to be included in our analyses, we required the individual to have at least two reads supporting the reference and alternative alleles, respectively, within a given tissue and at least five reads supporting each allele across the nine studied tissues. We also required the focal ASE site to be in Hardy-Weinberg equilibrium; determined using a chi-squared test with one degree of freedom and *α* = 0.005. These filters help reduce false signals of ASE resulting from genotyping errors at ASE sites.

For individual-gene pairs with multiple heterozygous sites that passed these filters, the site covered by the largest number of reads was analyzed.

Human imprinted genes as listed by *geneimprint* (downloaded from <http://www.geneimprint.com/site/genes-by-species>) were excluded from downstream analyses, as were highly polymorphic HLA genes (i.e. genes in the extended MHC region; bounded by SNPs rs498548 and rs2772390 plus 2Mb extensions on both sides).

At each individual-gene pair in each tissue that met our QC criteria, we calculated 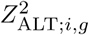 as described above. As an example, this resulted in 143,888 measurements of allelic imbalance at 18,735 unique loci in whole blood. We also calculated a combined-tissue ASE, wherein, within an individual, reads containing each allele at a focal ASE site were summed across tissues. This resulted in 372,473 measurements of combined-tissue allelic imbalance at 39,467 unique loci.

#### Contribution of cis-regulatory variants to ASE

To estimate the contributions of *cis*-regulatory variants to ASE, we determined the number of possible gene-regulatory variants at each locus. In each individual, we considered all heterozygous sites in *cis* to an ASE site to have potential effects on gene regulation.

We defined sites in *cis* to be those that lie within 10kb of the most upstream transcription start site of a gene containing an ASE site. We filtered sites not in Hardy-Weinberg equilibrium, determined using a chi-squared test with one degree of freedom and *α* = 0.005. We then counted, for each individual, the number of sites in *cis* to each ASE site that were called as heterozygous based on whole-genome sequencing data from the GTEx Project (21).

This resulted in 1,406,766 heterozygous sites in *cis* to 12,442 unique genes with measured ASE, with an average of approximately 100 *cis*-heterozygous sites per locus per individual. We then estimated the average contribution of a *cis*-heterozygous site to allelic imbalance using linear regression, as described in Eq. 13.

Due to correlation between data points (e.g. ASE was measured at the same gene in many individuals), we estimated 95% confidence intervals for the regression coefficient (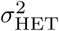) using a weighted jackknife as described in (34), excluding measurements from all individuals and tissues for a single gene in each sub-sample.

#### Variation in cis-regulation by allele frequency

To explore whether stabilizing selection acts on *cis*-regulatory variants in these ASE data, we tested whether there was a relationship between the allele frequency of a *cis*-heterozygous site and its contribution to allelic imbalance.

To do so, we binned *cis*-heterozygous sites by their minor allele frequency in the GTEx sample. We divided variants into singletons, doubletons, and spaced the remaining bins such they each contained approximately the same number of sites as the doubleton bin. The resulting bins have the following allele frequency cutoffs: (0.02, 0.04, 0.08, 0.15, 0.25, 0.37, 0.5).

For each ASE site in each individual, we counted the number of *cis*-heterozygous sites in each allele frequency bin. We then estimated the average contributions of *cis*-heterozygous sites in each bin to allelic imbalance using multiple regression, as described in Eq. 14.

To explore the confidence level of these estimates, we first permuted allelic imbalance measures across all individuals and genes. Second, we permuted allelic imbalance measures across all individuals within a gene. The first tests for global artifacts of the multiple regression the second tests for variation in allele frequency spectra and allelic imbalance across genes. In all cases, the total number of variants and the number of variants in each allele frequency bin were retained as in the original data. We performed 100 permutations for each of condition and compared the resulting estimates to those obtained in the original data.

### Data availability

RNA sequencing data for twins discordant for trisomy 21 were accessed from the Gene Expression Omnibus (GEO) data repository (accession GSE55426). Gene expression measurements (RPKM) across 13 tissues for healthy individuals with copy number variants were obtained from version 4 of the GTEx Project (dbGaP accession phs000424.v4.p1).

RNA sequencing and genotype data used in eQTL calling were accessed by application through the NIMH Center for Collaborative Genomic Studies on Mental Disorders. Instructions for requesting access to data can be found at https://www.nimhgenetics.org/access_data_biomaterial.php. Inquiries should reference the ‘Depression Genes and Networks study (D. Levinson, PI).’

1000 Genomes phase 3 data, used in polarizing genotypes to the human ancestral allele and in determining population allele frequencies are available from ftp://ftp.1000genomes.ebi.ac.uk/vol1/ftp/release/20130502/ALL.wgs.phase3_shapeit2_mvncall_integrated_v5b.20130502.sitez.vcf.gz. Details regarding the determination of human-chimp ancestral alleles are available in ftp://ftp.1000genomes.ebi.ac.uk/vol1/ftp/release/20130502/README_vcf_info_annotation.20141104.

Allele specific expression tables as well as genotype calls from whole genome sequencing from version 6p of the GTEx Project were accessed from dbGAP (accession phs000424.v6.p1).

## ACKNOWLEDGEMENTS

We thank the members of the Pritchard Lab for spirited conversations regarding this work. We also thank Lawrence Uricchio, Noah Zaitlen, and Ryan Hernandez for their comments. AH was funded in part by a fellowship from the Stanford Center for Computational, Evolutionary and Human Genomics (CEHG).

---

**Table S1.** Summary of data for analysis of the effects of winner’s curse and of limited statistical power on eQTL effect size estimates. To understand the impacts of winner’s curse, we called eQTLs in an ascertainment set and estimated their effect sizes in a validation set. To explore the impact of variable power to call eQTLs of the same effect across allele frequency bins, eQTLs were filtered to remove those with estimated effects that we would lack power to call at low allele frequencies. Here, we consider eQTLs with estimated effects larger than the minimum magnitude of estimated effect for significant, rare eQTLs (MAF<0.02). Especially rare eQTLs (MAF<0.02) are also excluded. ‘Power threshold’ shows this effect size cutoff for the different datasets.

|  | Full data | Ascertainment<br>*Validation* |
| --- | --- | --- |
| Sample size | 922 | 800<br>*122* |
| Number of eQTLs | 6,587 | 6,145 |
| Power threshold | 1.24×/ 0.81× | 1.27×/ 0.79× |
| Number of eQTLs<br>filtered for power | 2,702 | 2,333 |

**Fig. S1.**
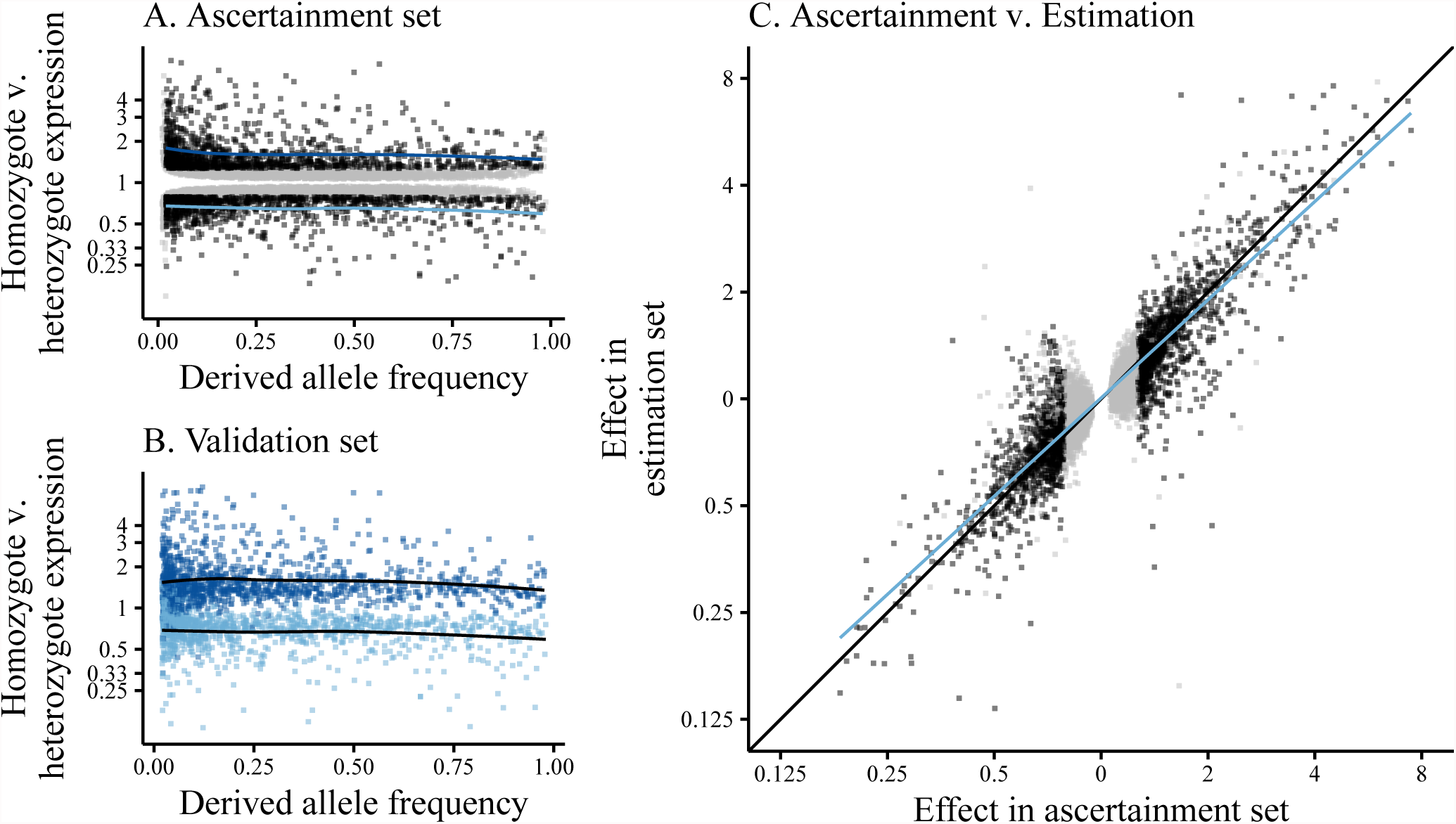
Comparison of eQTLs called using forward-stepwise mapping in an ascertainment set of 800 individuals, effect sizes estimated in 122 individuals. Data are whole blood RNA sequencing from the DGN cohort. *cis*-eQTLs were called in a 100kb window centered on the TSS of autosomal, protein-coding genes. Effect sizes are reported as the estimated expression of individuals heterozygous for a called eQTL relative to that of homozygous individuals and are polarized to the ancestral allele. (A) eQTL effect sizes (estimated in the ascertainment set) as a function of the derived allele frequency of the eQTL. eQTLs that we were uniformly powered to detect across allele frequencies (estimated effect greater than the minimum effect size estimated for eQTLs with MAF<0.02) shown in black, others (including especially rare eQTLs, MAF<0.02) in grey. Loess-fits of derived allele frequency vs eQTL effect size (for well-powered eQTLs, black points, for expression-increasing and -decreasing eQTLs, respectively) are shown in blue. (B) eQTL effect sizes (estimated in the validation set) as a function of the derived allele frequency of the eQTL. Only eQTLs that we were well-powered to detect across allele frequencies and with MAF>0.02 are plotted. Points are colored by the direction of the eQTL effect in the ascertainment set; expression-increasing eQTLs (derived allele predicted to increase expression) shown in dark blue, expression-decreasing eQTLs in light blue. Loess-fits of derived allele frequency vs eQTL effect size (for expression-increasing and -decreasing eQTLs, respectively) are shown in black. (C) Correlation between effect sizes estimated in the larger, ascertainment sample (x-axis) and the smaller, validation sample (y-axis). eQTLs that we were uniformly powered to detect across allele frequencies (estimated effect greater than the minimum effect size estimated for eQTLs with MAF<0.02) shown in black, others (including rare eQTLs, MAF<0.02) in grey. *y* = *x* line is shown in black, least-squares regression line shown in blue.

**Fig. S2.**
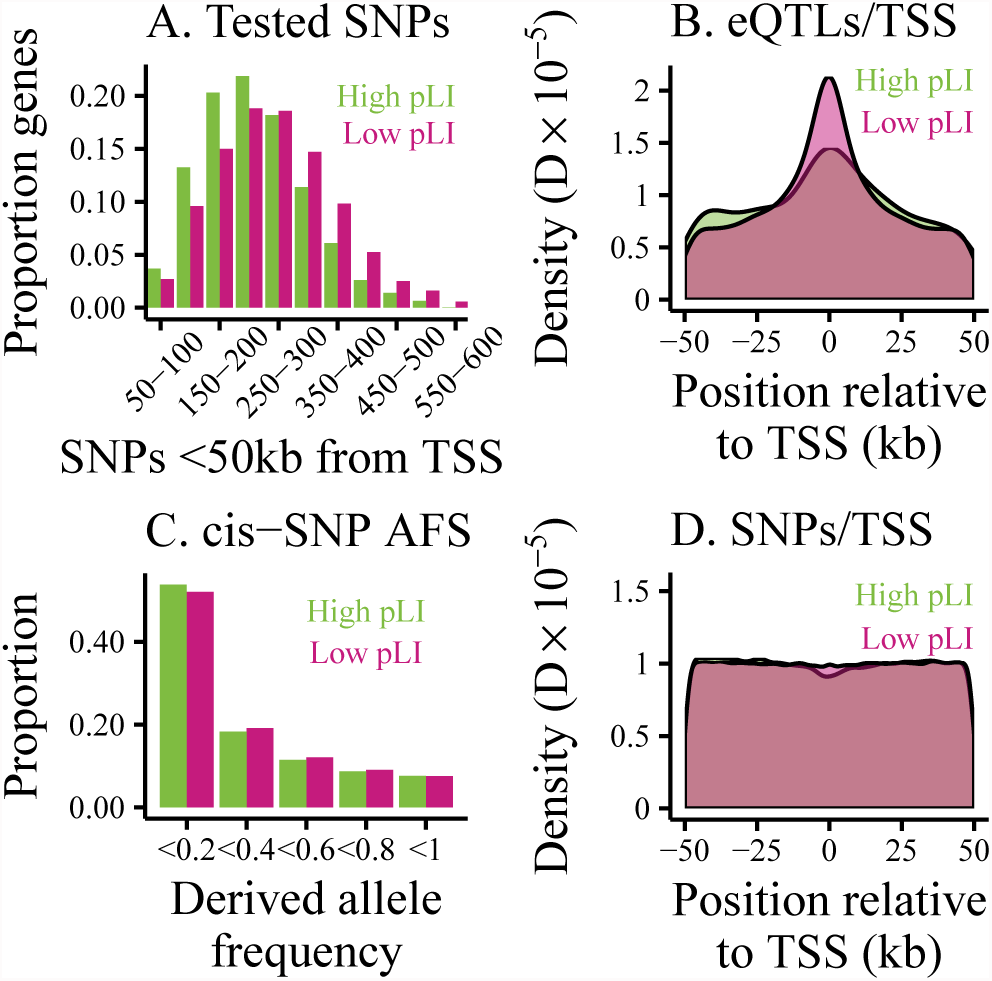
Profile of candidate eQTLs (SNPs with MAF>0.05 within 50kb of the TSS of an autosomal, protein-coding gene) around high- and low-*pLI* genes. (A) and (C) compare the numbers and allele frequencies of SNPs around high- and low-*pLI* genes, while (B) and (D) compare the distribution of called eQTLs and SNPs around transcription start sites. (A) Histogram of the number of *cis*-SNPs tested as eQTLs per gene. (C) Histogram of the derived allele frequencies of *cis*-SNPs. (B) Density of called eQTLs relative to the TSS. (D) Density of candidate eQTLs relative to the TSS.

**Fig. S3.**
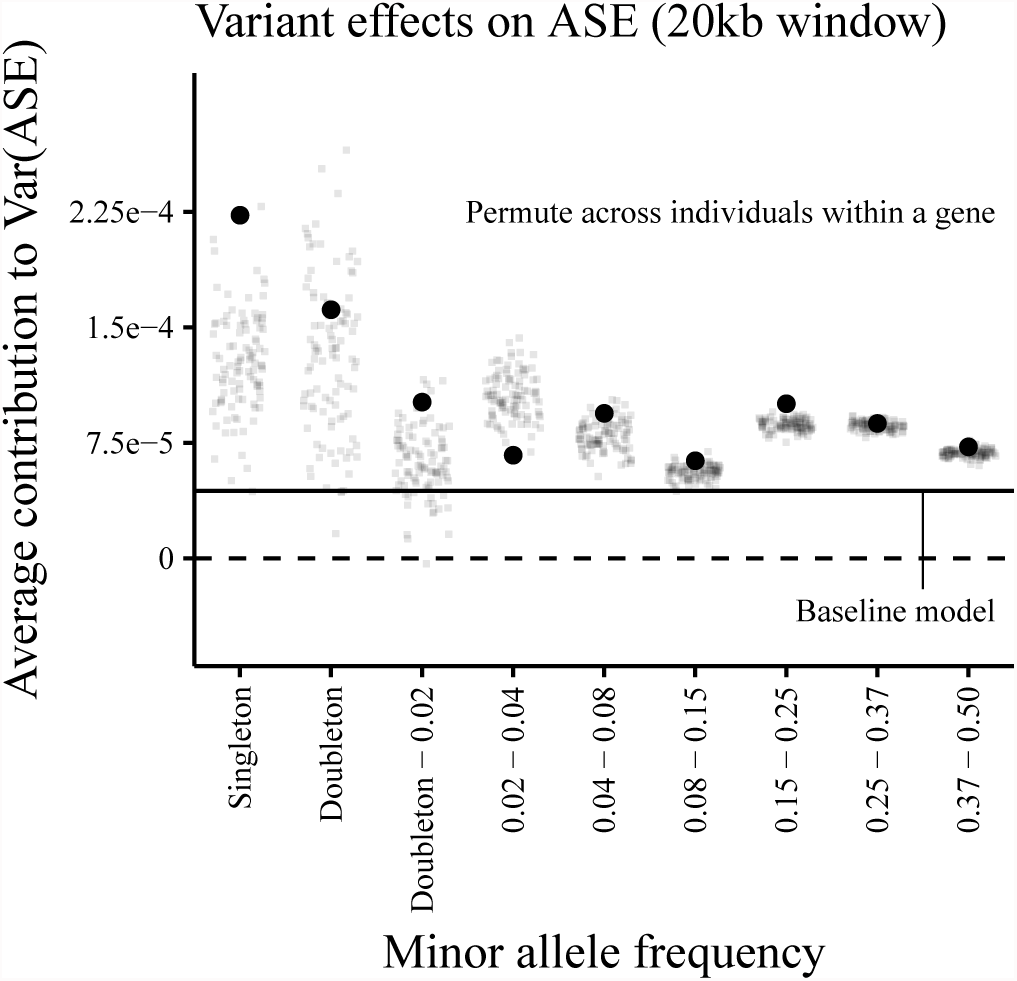
Effects of *cis*-genetic variants (those within 10kb of the TSS) on allelic imbalance. Estimates from combined-tissue ASE (reads spanning each ASE site in a single individual are summed, regardless of the tissue in which they were sampled) are shown as large points. Small points represent estimates from 100 permutations of ASE measurements across individuals within a gene. The estimated singleton effect size from real data is significantly greater than those from the permutations. However, correlation in the allele frequency spectra at a locus across individuals clearly contributes to our effect size estimates. One interpretation of this is that genes vary in their level of constraint on expression. This could lead to subsets of genes with different frequency spectra and different ASE profiles.

**Fig. S4.**
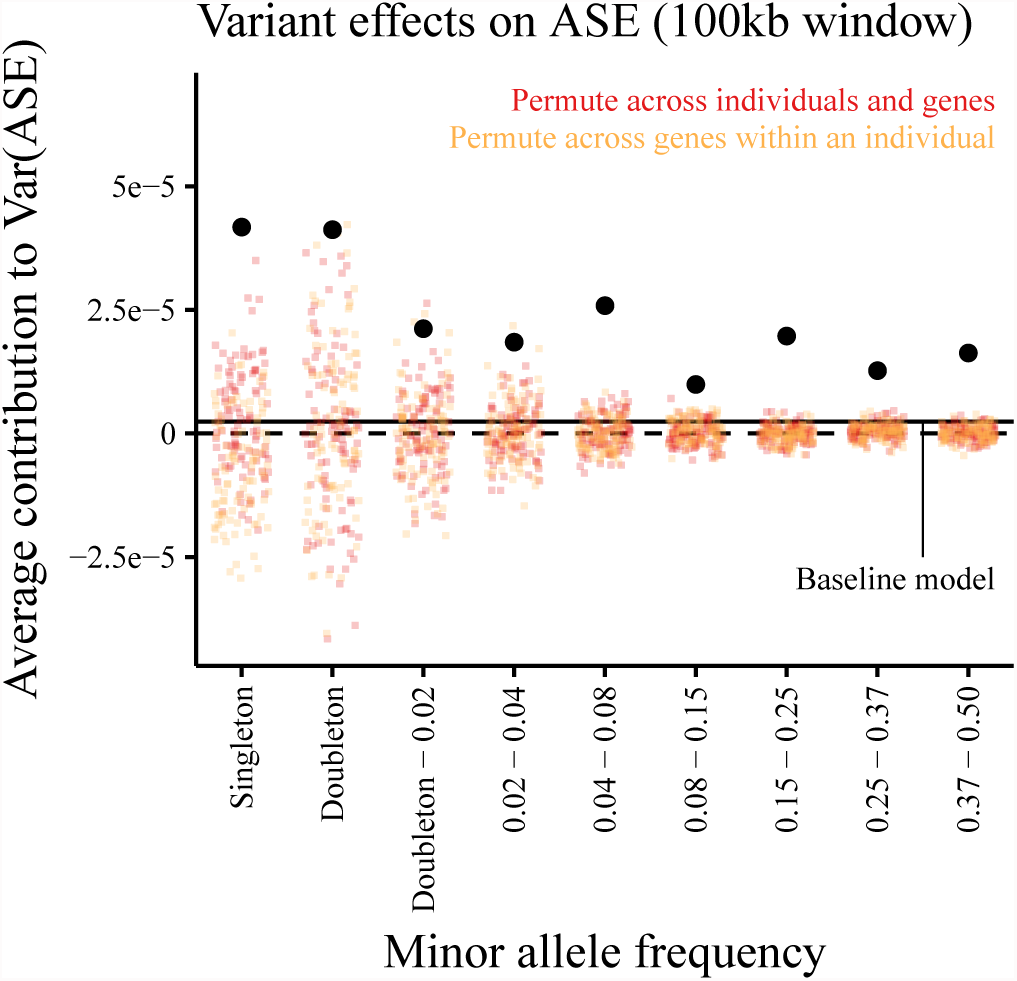
Effects of *cis*-genetic variants (those within 50kb of the TSS) on allelic imbalance. Estimates from combined-tissue ASE (reads spanning each ASE site in a single individual are summed, regardless of the tissue in which they were sampled) are shown in black. Colored points represent estimates from 100 permutations of (1) ASE measurements across genes and individuals (red) and (2) ASE measurements across genes within an individual (orange). Horizontal lines mark zero effect (dashed) and the average variant effect estimated by regressing allelic imbalance on the total number of *cis*-heterozygous sites (solid).

**Fig. S5.**
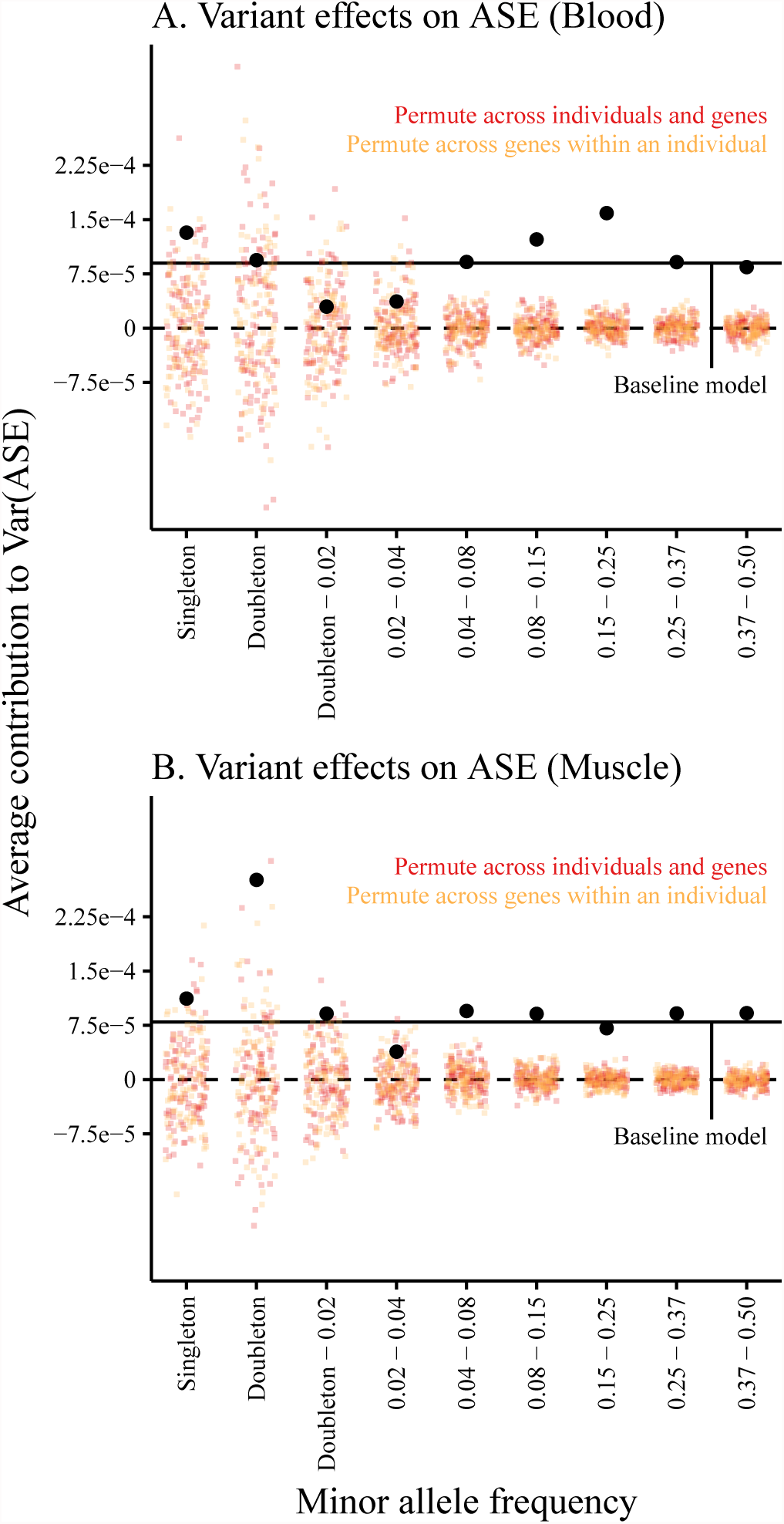
Effects of *cis*-genetic variants (those within 10kb of the TSS) on allelic imbalance in individual tissues. Estimates from multiple regression shown in black. Colored points represent estimates from 100 permutations of (1) ASE measurements across genes and individuals (red) and (2) ASE measurements across genes within an individual (orange). Horizontal lines mark zero effect (dashed) and the average variant effect estimated by regressing allelic imbalance on the total number of *cis*-heterozygous sites (solid). (A) Estimates and permutations based on all individual-gene pairs sampled in whole blood. (B) Estimates and permutations based on all individual-gene pairs sampled in skeletal muscle.

**Fig. S6.**
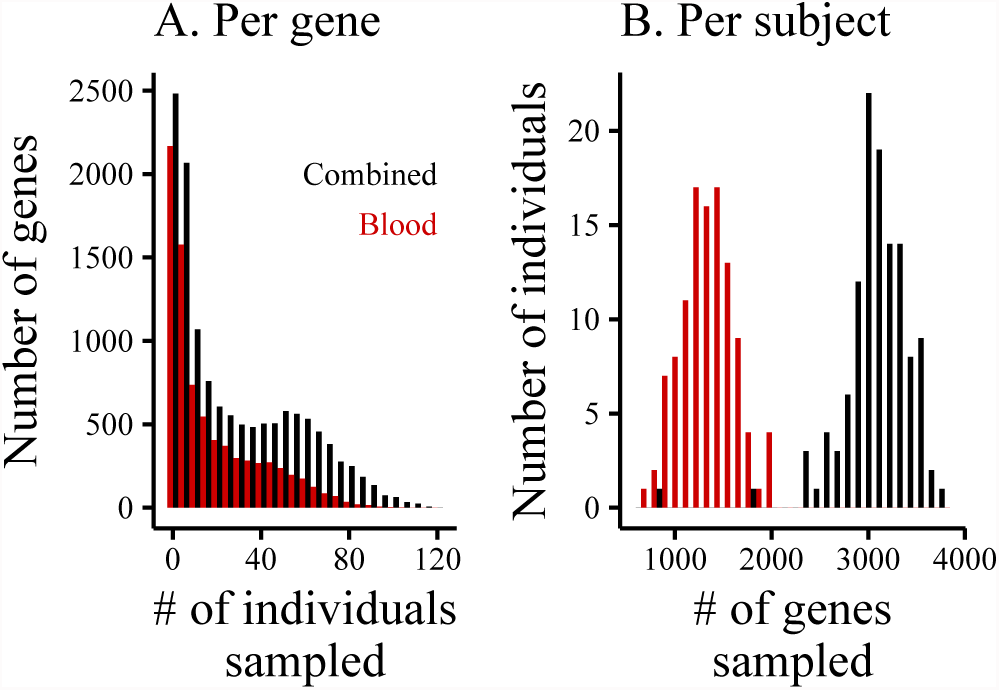
Summary of ASE sample sizes across tissues. Data from version 6 of the GTEx Project. ‘Combined’ (shown in black) refers to combined-tissue ASE (reads spanning each ASE site in a single individual are summed, regardless of the tissue in which they were sampled). The nine best-sampled GTEx tissues are included. ‘Blood’ (shown in red) are ASE measurements from whole blood alone. (A) Histogram of the number of individuals with measured allelic imbalance per gene. (B) Histogram of the number of genes with measured allelic imbalance per individual.

**Fig. S7.**
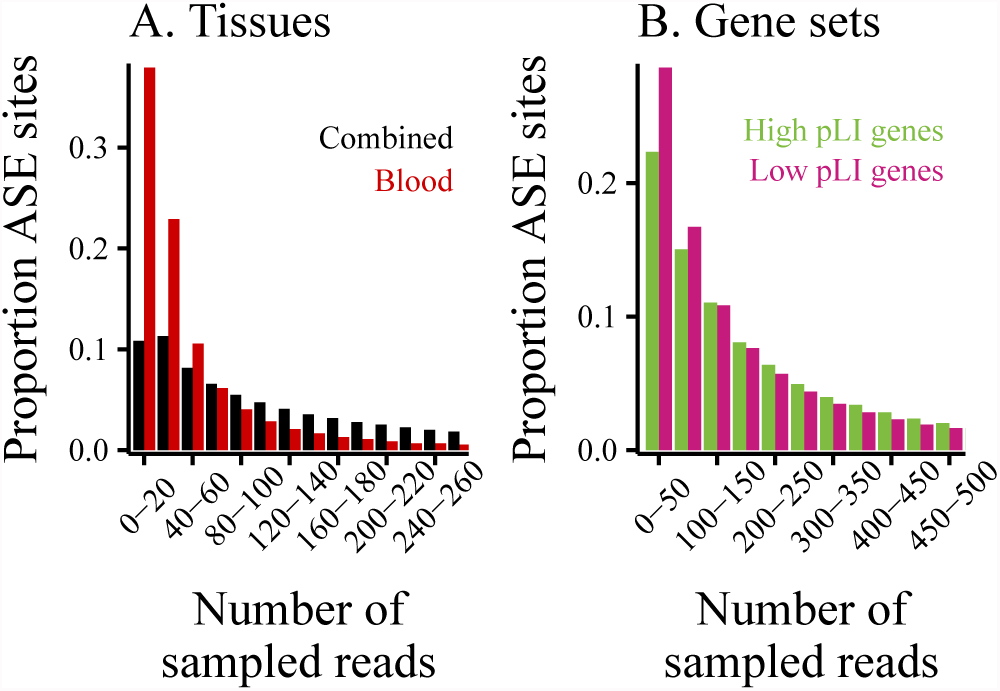
Summary of ASE read depth across tissues and gene sets. Data from version 6 of the GTEx Project. (A) Histogram of read depth across tissues. ‘Combined’ (shown in black) refers to combined-tissue ASE (reads spanning each ASE site in a single individual are summed, regardless of the tissue in which they were sampled). The nine best-sampled GTEx tissues are included. ‘Blood’ (shown in red) are ASE measurements from whole blood alone. (B) Histogram of read depth for combined-tissue ASE across gene sets (from Lek et al. 13).

**Fig. S8.**
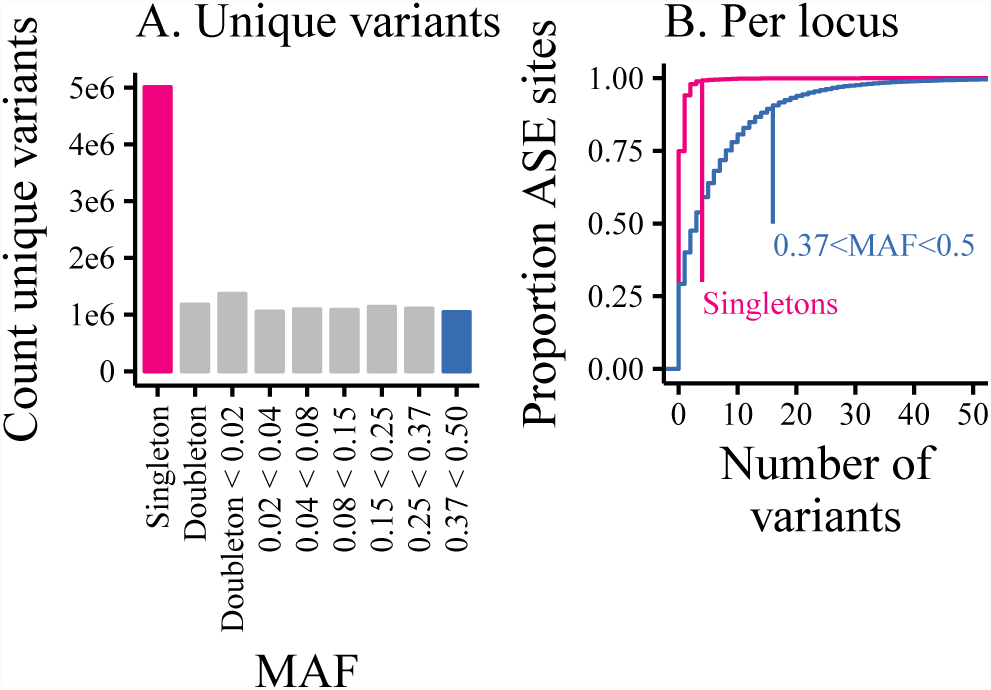
Summary of allele frequencies of variants (heterozygous in at least one individual) within 10kb of the TSS of genes with measured ASE in version 6 of the GTEx Project. (A) Histogram of the count of unique variants in each allele frequency bin. (B) Cumulative distribution of the number of singletons (pink) and the most common variants (0.37<MAF<0.5, blue) per individual-gene pair.

